# Retrograde mitochondrial transport is required for mitochondrial biogenesis in zebrafish neurons

**DOI:** 10.1101/2025.09.29.679307

**Authors:** Angelica E Lang, Chris Stein, Roger Schultz, Catherine M Drerup

## Abstract

To maintain a functional mitochondrial population in a long-lived cell like a neuron, mitochondria must be continuously replenished through the process of mitochondrial biogenesis. Because most mitochondrial proteins are nuclear encoded, mitochondrial biogenesis requires communication between mitochondria and the nucleus. This can be a challenge in a large, compartmentalized cell like a neuron in which a large portion of the mitochondrial population is in neuronal compartments far from the nucleus. Using in vivo assessments of mitochondrial biogenesis in zebrafish neurons, we determined that mitochondrial transport between distal axonal compartments and the cell body is required for sustained mitochondrial biogenesis. Estrogen-related receptor transcriptional activation links transport with nuclear expression of mitochondrial genes. Together, our data support a role for retrograde feedback between axonal mitochondria and the nucleus for regulation of mitochondrial biogenesis in neurons.

## Introduction

Mitochondria are critical for neuronal function and survival. These organelles have many essential roles, including production of ATP, calcium buffering, and the production of metabolites that can function as signaling molecules ^1^. For a neuron to maintain a functional mitochondrial population, mitochondria must be replenished through the process of mitochondrial biogenesis ^2^. In other cell types, such as myocytes and adipocytes, indicators of mitochondrial health and function like AMP/ATP and NAD+/NADH ratios as well as calcium signaling stimulate the transcriptional control of mitochondrial biogenesis in the nucleus ^3–6^. These upstream signals can activate transcriptional regulators like the PGC-1 (Peroxisome Proliferator-activated Receptor Gamma Coactivator 1) family of transcriptional coactivators ^7^. PGC-1α, considered the master regulator of mitochondrial biogenesis, can be activated by phosphorylation or de-acetylation by AMP-activated protein kinase (AMPK) or Sirtuin 1 (SIRT1) respectively ^3,4,6^. AMPK activity is stimulated by increases in the AMP/ATP ratio while SIRT1 activity depends on local NAD+ levels. Once activated, PGC-1α can induce the expression of nuclear-encoded mitochondrial proteins through the activation of transcription factors like the nuclear respiratory factors (NRF1/2) or the estrogen related receptors (ERRα/β/γ) ^7–10^. How neurons, with mitochondrial populations spread throughout dendritic and axonal arbors, oftentimes large distances from the nucleus, sense mitochondrial health and function to activate these transcriptional pathways is an open and critical question ^2^.

Mitochondrial biogenesis requires the division and expansion of existing organelles to increase mitochondrial mass. This requires the coordination of several cellular processes. Mitochondrial DNA (mtDNA) must be replicated and the mtDNA nucleoids segregated, after which mitochondrial fission occurs at or near the middle of the organelle ^11,12^. Daughter mitochondria are expanded through transcription, translation, and mitochondrial import of nuclear encoded gene products as well as cellular and mitochondrial derived lipids ^2,13^. Mitochondria also synthesize proteins encoded by mtDNA (thirteen in vertebrates) in the mitochondrial matrix. Given the dependence on coordinated mitochondrial and nuclear gene expression, it is unsurprising that a substantial amount of mitochondrial biogenesis occurs in the perinuclear region in non-neuronal cells ^14,15^. In neurons, evidence of mitochondrial biogenesis, including mtDNA replication and synthesis of mitochondrial proteins, has been observed in both the neuronal cell body and in axons of chick, rat, and drosophila ^13,16–19^. Local mitochondrial biogenesis is thought to support short-term, local changes in energy need at the synapse ^16,19,20^. The respective contributions of cell body-based verses local mitochondrial biogenesis for support of synaptic mitochondrial populations is still a subject of investigation. For both, however, nuclear expression of mitochondrial genes would be required to produce the >1000 proteins necessary for functional organelles.

Mitochondria are actively transported between the cell body and axon terminal by microtubule-based transport. In neuronal axons, kinesin motor proteins are responsible for axon terminal directed (anterograde) mitochondrial transport while the Cytoplasmic dynein motor transports mitochondria from the periphery back to the cell body (retrograde) ^21–24^. A complement of proteins including Trak1/2 and Miro1/2 have been shown to scaffold mitochondria to these motors for transport ^25–31^. Disrupting mitochondrial transport into the axon impairs axon development and regeneration after injury ^32–34^. Inhibition of retrograde mitochondrial transport from neurites back to the cell body causes mitochondrial accumulation in axonal and dendritic processes and impaired mitochondrial health at the synapse ^35–37^. However, the role of retrograde mitochondrial transport in mitochondrial population maintenance in neurons remained undefined.

We hypothesized that transport of mitochondria from distal regions of the neuron to the cell body could link mitochondrial population surveillance and regulation of mitochondrial biogenesis frequency in neurons. Using zebrafish neurons, we show that reducing mitochondrial retrograde transport from the axon terminal back to the cell body reduces markers of mitochondrial biogenesis, including mtDNA replication, nuclear transcription of mitochondrial genes, and the production of mitochondrial biomass in the cell body. mtDNA replication is also reduced in the axon terminal, suggesting both local and cell body-based mitochondrial biogenesis may be disrupted. Analysis of mitochondrial turnover at the synapse revealed that reduced mitochondrial biogenesis correlates with reductions in cell body-derived mitochondrial material transported to the axon terminal. Widespread disruption to mitochondrial gene expression in neurons lacking mitochondrial retrograde transport suggests transport plays an important role in regulation of transcriptional programs necessary for mitochondrial biogenesis. We present evidence that this crosstalk is mediated by NAD+/SIRT1-mediated deacetylation of Estrogen related receptors (ERR) which regulates ERR-dependent transcriptional activity. Together, our work supports a model in which the retrograde transport of mitochondria is a powerful signaling mechanism to regulate mitochondrial biogenesis in neurons.

## Results

### Inhibiting mitochondrial retrograde transport disrupts markers of mitochondrial biogenesis in neuronal cell bodies

We used larval zebrafish posterior lateral line (pLL) sensory neurons to study the dynamics of mitochondrial biogenesis in vivo. The pLL system is composed of afferent neurons and sensory organs called neuromasts situated along the trunk ^38^. The cell bodies of pLL neurons are clustered in a ganglion (pLLg) near the head while the afferent axons extend down the body and branch into axon terminals that innervate neuromasts (Fig. 1a). Axon extension and synapse formation occurs by 4 days post fertilization (dpf; ^39^), allowing us to use this system to study mitochondrial populations in a fully formed neuronal circuit in the translucent zebrafish larva. To disrupt retrograde mitochondrial transport in pLL neurons, we used an existing mutant line with a unique loss of function mutation in the gene encoding Actr10 (*actr10^nl^*^15^; hereafter referred to as *actr10;* ^35,37^). Actr10 is a component of the Cytoplasmic dynein accessory complex dynactin. Complete loss of Actr10 function results in failed cell division, likely due to loss of all dynein motor function ^40^. In contrast, the *actr10^nl^*^15^ mutation in zebrafish causes loss of mitochondrial interaction with the dynein motor, leading to loss of retrograde (cell body directed) mitochondrial transport and accumulation of mitochondria in axon terminals (^35^; Fig. 1b, c). This disruption is specific to mitochondria, as other retrogradely moving cargos, including late endosomes, lysosomes, and signaling molecules, are transported normally. Additionally, transmission electron microscopy (TEM) analysis showed accumulations of only mitochondria in axon terminals ^35^. Intriguingly, in addition to mitochondrial accumulation in the distal axons, we found a severe reduction in mitochondrial density (mitochondrial area/cytosolic area) in the cell body of *actr10* neurons (Fig. 1b, c). We asked if this loss of mitochondrial density could be due to loss of mitochondrial biogenesis in *actr10* mutants.

**Figure 1:**
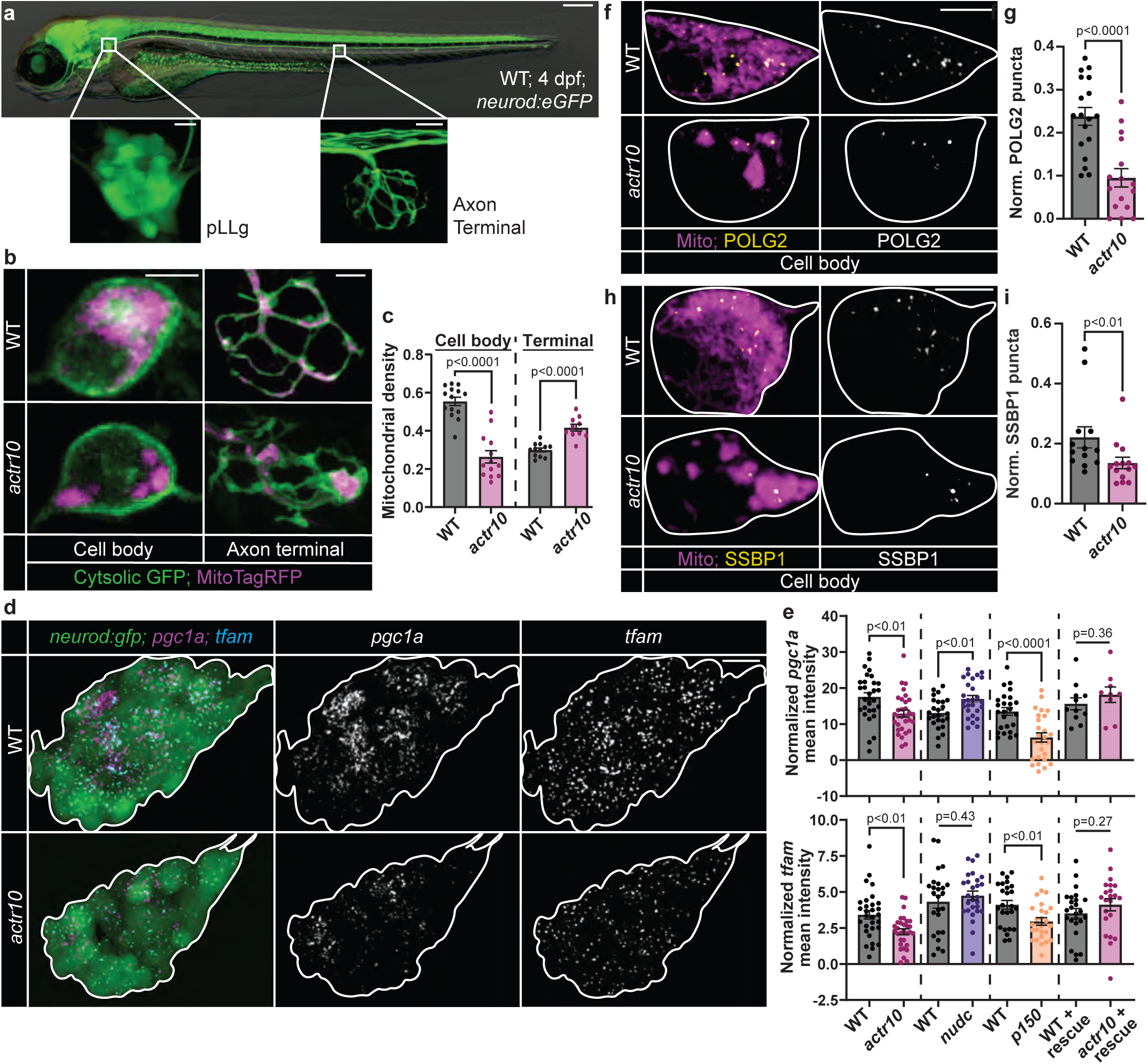
Mitochondrial cell body density and biogenesis markers are reduced in *actr10* mutants. (a) 4 dpf zebrafish transgenic larva carrying the *TgBAC(neurod:egfp)^nl^*^1^ transgene which labels neurons with GFP. The posterior lateral line gangion (pLLg) and a pLL axon terminal are boxed and magnified. (b) 4 dpf pLLg cell body and pLL axon terminal expressing cytosolic GFP and mitochondria-localized TagRFP (MitoTagRFP, magenta) in wild type and *actr10* mutants. (c) Quantification of mitochondrial density (mitochondrial area/cytosolic area; ANOVAs). (d) HCR RNA FISH labeling of *pgc1a* and *tfam* mRNA in the pLLg of wild type and *actr10* mutants carrying the *TgBAC(neurod:egfp)^nl^*^1^ transgene (white outline). (e) pLLg mean fluorescence intensity of *pgc1a* and *tfam* normalized to background (ANOVAs) in wild type, *actr10*, *nudc*, *p150*, and wild type/*actr10* larvae expressing Actr10 in neurons (+ rescue). (f) pLL cell body (outlined) expressing POLG2-GFP and mitoTagRFP (Mito, Magenta). (h) SSBP1 immunostaining in pLL cell body mitochondria, visualized with mitochondria-localized GFP (Mito, magenta; cell outlined). (g, i) Number of POLG2 or SSBP1 puncta normalized to mitochondrial volume (Wilcoxon). Scale bars: (a) full larva = 200 μm, insets = 10 μm; (b, f, & h) = 5 μm; (d) = 10 μm. All data are mean ± SEM and points represent individual larvae.

Mitochondrial biogenesis requires coordinated transcription of nuclear encoded mitochondrial genes. This is largely controlled by the PGC-1 family of transcriptional coactivators and their associated transcription factors, including NRF-1/2. Downstream of this transcriptional activation is a plethora of nuclear-encoded mitochondrial genes, including *tfam* ^7,10^. We first assessed the expression of *pgc1a*, *pgc1b*, *nrf1*, and *tfam* using colorimetric in situ hybridization and found that all were noticeably reduced in *actr10* mutants (Supplementary Fig. 1a, b). To quantify gene expression of biogenesis markers in the pLLg, we turned to hybridization chain reaction RNA fluorescence in situ hybridization (HCR RNA FISH; ^41^). We assessed *pgc1a* and *tfam* expression and confirmed both were significantly reduced in *actr10* mutants (Fig. 1d, e). Neuronal rescue of Actr10 in a stable transgenic line (*Tg(5kbneurod:mRFP-actr10)^nl^*^22^; ^35^) rescued neuronal mitochondrial density (Supplementary Fig. 1c, d) as well as *pgc1a* and *tfam* expression (Fig. 1e) confirming loss of Actr10 in neurons underlies this phenotype. To determine if the changes to gene expression in *actr10* mutants could be explained by general disruptions to retrograde cargo transport, we quantified gene expression in control mutant strains with loss of function mutations in *p150a/b* (duplicated gene in teleost fish requiring double knockout) and *nudc* ^35,42^. p150 is a component of the dynactin complex that, when mutated, disrupts all retrograde cargo transport, including mitochondria ^35,43–45^. NudC is a dynein regulator that, when mutated in zebrafish neurons, disrupts the retrograde transport of late endosomes and autophagosomes without altering the localization or transport of mitochondria ^42^. *pgc1a* and *tfam* expression were reduced in *p150* mutants but unaffected in *nudc* mutants, further suggesting the decreased expression of these mitochondrial biogenesis markers was due to the loss of retrograde mitochondrial transport (Fig. 1e).

We next analyzed a critical step during biogenesis, mtDNA replication, using POLG2-GFP, a fluorescently tagged version of the processivity subunit of the mitochondrial DNA polymerase. POLG2-GFP has been previously shown to specifically localize to mtDNA nucleoids actively undergoing replication ^12,46^. We expressed POLG2-GFP in pLL neurons and quantified the number of puncta, indicative of replicating mtDNA nucleoids, relative to mitochondrial volume. The number of POLG2 puncta was significantly reduced in *actr10* pLL cell body mitochondria, indicative of reduced mtDNA replication (Fig. 1f, g). To confirm these results, we performed immunolabeling for single stranded binding protein 1 (SSBP1), which transiently associates with mtDNA to facilitate replication ^47^. The number of SSBP1 puncta relative to mitochondrial volume was significantly reduced in *actr10* pLL cell bodies (Fig. 1h, i). This phenotype is not due to loss of mtDNA as the density of nucleoids labelled with the mtDNA nucleoid protein TFAM (transcription factor A, mitochondria; ^48^) is not decreased in *actr10* cell body mitochondria (Supplementary Fig.1e, f). As expected, due to the reduced mitochondrial density in the cell body and in line with loss of *tfam* mRNA based on HCR RNA FISH quantification, overall number of TFAM puncta are reduced in *actr10* mutant cell bodies (Supplementary Fig. 1g). mtDNA nucleoid size was unchanged between wild type and *actr10* mutants (Supplementary Fig. 1h). Together with our previous results demonstrating no deficits in measures of mitochondrial health and function in *actr10* mutant pLL neuronal cell bodies^37^, this suggests decreases in mtDNA replication are not due to mitochondrial stress in the cell body. Mitochondrial biogenesis has also been demonstrated to occur in the distal axon through assessment of mtDNA replication ^16,19,20^. To determine if local mitochondrial biogenesis was also affected in *actr10* mutants, we quantified mtDNA replication in the pLL axon terminal. Both POLG2 and SSBP1 puncta were significantly reduced in *actr10* mutants, suggesting both cell body and axon terminal mitochondrial biogenesis are disrupted by loss of retrograde mitochondrial transport (Supplementary Fig. 1i-l).

### Nuclear transcription of mitochondrial genes is disrupted in actr10 mutant neurons

Given the changes to mtDNA replication and transcription of mitochondrial biogenesis related genes, we decided to look more broadly at how nuclear gene expression of mitochondrial proteins may be altered in *actr10* mutants. We isolated pLL neurons using fluorescence activated cell sorting (FACS) from a novel transgenic line (*Tg(hsp70:eGFP-SILL*)*^uwd12Tg^*; Supplementary Fig. 2a, b) which expresses GFP in pLL neurons. FACS gating parameters were optimized to sort true GFP-expressing cells and exclude autofluorescence particularly apparent in the yolk (Supplementary Fig. c). We then performed bulk RNA-sequencing on FACS sorted cells to analyze the transcriptomes of wild type and *actr10* mutant neurons (Fig. 2a). Differential gene expression analysis and cross comparisons to the MitoCarta database confirmed that mitochondrial genes are significantly downregulated in *actr10* mutants compared to wild type siblings (Fig. 2b, c; ^49^). In contrast, there were only 3 mitochondrial genes significantly upregulated (p=0.999, Fisher’s exact test). Gene Set Enrichment Analysis (GSEA) of Gene Ontology terms identified Mitochondrial Protein Complexes to be the most significantly de-enriched term in the dataset (Fig. 2d and Supplementary Fig. 2d) and KEGG pathway enrichment analysis showed strong enrichment for mitochondrial pathways in downregulated genes (Supplementary Fig. 2e). To confirm our RNA-seq results, we used HCR RNA FISH to quantify pLL ganglion expression of ten downregulated genes and two genes that were not altered in our dataset. These analyses aligned with our RNA-seq data, confirming loss of these mitochondrial transcripts in *actr10* mutants (Fig. 2e-h). Additionally, we confirmed these results using colorimetric in situ hybridization of a subset of these genes, showing reduced expression of *atp5md* and *cox5ab* but not the controls *polb* and *tomm40* (Supplementary Fig. 2f, g). Finally, to assess the specificity of the transcriptomic changes, we included positive and negative control mutant strains *p150* and *nudc* described above ^35,42^. As predicted, *p150* but not *nudc* mutants have a loss of *cox5ab* expression (Fig. 2e, f). Together, this transcriptomic analysis suggests that mitochondrial retrograde transport modulates nuclear expression of a subset of mitochondrial genes.

**Figure 2:**
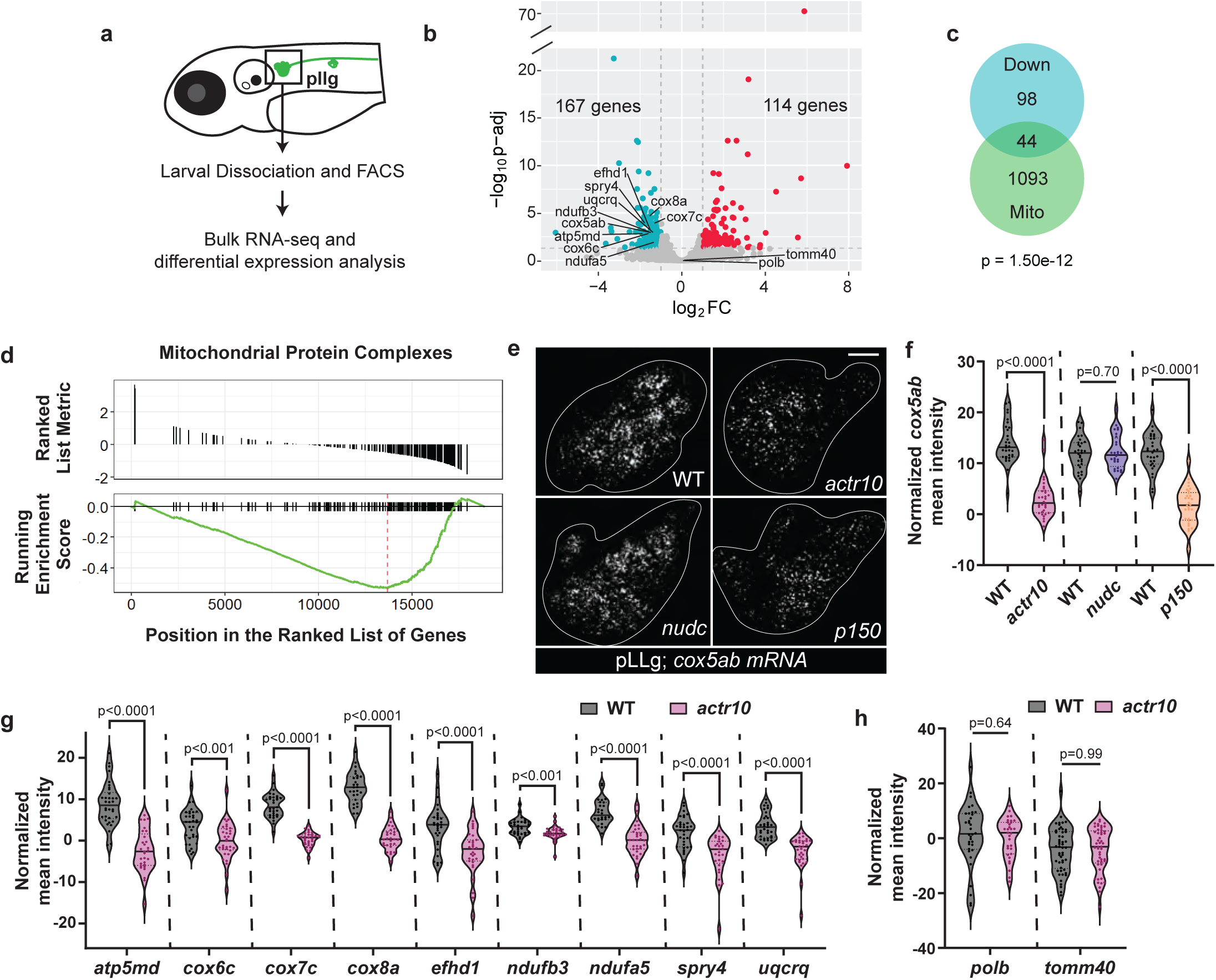
Mitochondrial gene expression is downregulated in *actr10* mutants. (a) RNA-sequencing strategy. (b) Volcano plot showing differential gene expression between wild type and *actr10* mutants. The plot shows -log10(adjusted p-value) vs. log2(Fold Change). Blue dots represent genes significantly downregulated while red dots represent genes significantly upregulated in *actr10* mutants. Genes selected for secondary analysis using HCR RNA FISH are labeled. (c) Overlap between downregulated genes and the MitoCarta3.0 database of the mitochondrial proteome (Fisher’s exact test). Human homologs of downregulated zebrafish genes were used for analysis. Duplicated zebrafish paralogues were only counted once. (d) GSEA plot for Mitochondrial Protein Complexes, the most significantly de-enriched gene ontology term in *actr10* mutants relative to wild type. (e) Representative images of HCR RNA FISH labeling of *cox5ab* mRNA in the pLLg (outlined in white) for wild type, *actr10*, *nudc*, and *p150* mutants. Scale bar = 10 μm. (f) pLLg mean fluorescence intensity of *cox5ab* normalized to background (ANOVAs). (g) pLLg mean fluorescence intensity normalized to background for downregulated mitochondrial genes in wild type vs. *actr10* mutants (ANOVAs). (h) pLLg mean fluorescence intensity normalized to background for control genes in wild type vs. *actr10* mutants (ANOVAs). (f-h) Data plot lines represent the median and quartiles; data points represent individual larvae.

### Impaired mitochondrial biogenesis decreases the contribution of cell body-derived mitochondria to the axon terminal

We next wanted to determine if these changes in measures of mitochondrial biogenesis translated into a decrease in the production of new organelles. In neurons, we predicted that at least a subset of mitochondria produced by biogenesis in the cell body would be transported in the anterograde direction to support the population of mitochondria in the axon terminal ^2^. To test this, we used photo-labeling strategies to assess the rate of cell body-derived mitochondrial addition to the axon terminal in wild type animals. We expressed photoactivatable GFP (PA-GFP) localized to the mitochondrial matrix in pLL neurons and locally activated the GFP only in the cell body. Then, we quantified the addition of cell body derived mitochondrial material to the axon terminal at multiple time-points after photo-activation. We found that by 4 to 8 hours post-activation (hpa), ∼50% of the mitochondrial density in the axon terminal contained material that was cell body-derived (Supplementary Fig. 3a-d). Next, we asked if the amount of cell body-derived mitochondrial material transported to the axon terminal correlated with turnover of the axon terminal mitochondrial population. Published data has shown that axon terminal mitochondria are consistently lost over a period of hours in pLL axon terminals ^37^. Using precise activation of matrix-localized PA-GFP in axon terminal mitochondria followed by post-activation imaging, we determined that ∼50% of the labeled mitochondrial density is absent from the axon terminal by 8 hpa (Supplementary Fig. 3e-g). This complements the estimated rate of mitochondrial entrance into the axon terminal from the cell body and suggests that cell body-derived mitochondria may play a significant role in maintenance of the axon terminal mitochondrial population.

Given the role for cell body-derived mitochondria in axon terminal turnover, we predicted that decreased biogenesis in the cell body would result in a progressive loss of mitochondrial addition to the axon terminal. To test this, we assessed changes to mitochondrial biogenesis and transport in *actr10* mutants across multiple timepoints post-axon extension (2, 3, and 4 dpf). Initial axon extension of primary pLL axons is complete by 2 dpf with functional synapses apparent by 4 dpf ^38,39,50^. Assessment of mitochondrial density in axon terminals and cell bodies of *actr10* mutants demonstrated that mitochondrial density was elevated in axon terminals as early as 2 dpf (Supplementary Fig. 4a, b) and cell body mitochondrial density was notably depleted by 3 dpf (Supplementary Fig. 4c, d). Quantification of the biogenesis markers *pgc1a* and *tfam* using HCR RNA FISH demonstrated that loss of cell body mitochondrial density at 3 dpf corresponded with loss of *pgc1a* expression, followed by loss of *tfam* expression by 4 dpf (Fig. 1d, e and Supplementary Fig. 4e, f). This time course confirms that loss of cell body mitochondrial density correlates with decreased markers of mitochondrial biogenesis.

We predicted that the progressive loss of cell body mitochondrial biogenesis in *actr10* mutants would result in reduced anterograde mitochondrial transport. Live imaging of mitochondrial movement in pLL axons demonstrated that retrograde transport was disrupted at 3 dpf while anterograde transport frequency was not altered until 4 dpf (Supplementary Fig. 4g, h and Supplementary Movies 1-4). This time series demonstrated retrograde mitochondrial transport is lost first in *actr10* mutants, leading to mitochondrial accumulation in the axon terminal. Only after retrograde mitochondrial transport is inhibited does mitochondrial biogenesis fail, leading to loss of cell body mitochondria and reduced anterograde transport of mitochondria towards axon terminals (summarized in Supplementary Fig. 4i).

Together, our data suggest that anterograde mitochondrial transport supplies the axon terminal with cell body-derived mitochondria produced through biogenesis. To test this, we directly stimulated mitochondrial biogenesis and analyzed axon terminal mitochondrial addition. To increase mitochondrial biogenesis, we used a stable transgenic zebrafish line overexpressing human PGC-1α in neurons (*Tg(-5kbneurod1:PGC1α-2A-mRFP)^uwd9Tg^*; ^51^). PGC-1α overexpression in this line increased *tfam* expression in pLL neurons and has been previously shown to increase mitochondrial density in the axon terminal, confirming increased mitochondrial biogenesis (Supplementary Fig. 4j; ^51^). To determine if increased mitochondrial biogenesis increases the addition of cell body-derived mitochondria to the axon terminal, we used photoconversion and mitochondrial tracking. For this, we used a dual transgenic line expressing PGC-1α (*Tg(-5kbneurod1:PGC1α-2A-mRFP)^uwd9Tg^*) and mEos, a photoconvertible protein, localized to the mitochondrial matrix (*Tg(-5kbneurod1:mito-mEos)^y^*^586^*^Tg^*; ^37,52^). We photoconverted the mitochondrial population in the pLL ganglion at 4 dpf and imaged axon terminals 4 hours post-conversion (hpc; Fig. 3a). By 4 hpc, cell body-derived mitochondrial density in the axon terminal was significantly increased in PGC-1α transgenics, confirming that mitochondrial biogenesis in the cell body impacts mitochondrial addition to the axon terminal population (Fig. 3b, c). Next, we assessed the density of cell body-derived mitochondria in *actr10* mutant axon terminals using the same photoconversion strategy. This demonstrated a 50% reduction in the contribution of cell body-derived mitochondria to the axon terminal population in *actr10* mutants compared to wild type (Fig. 3b, c).

**Figure 3:**
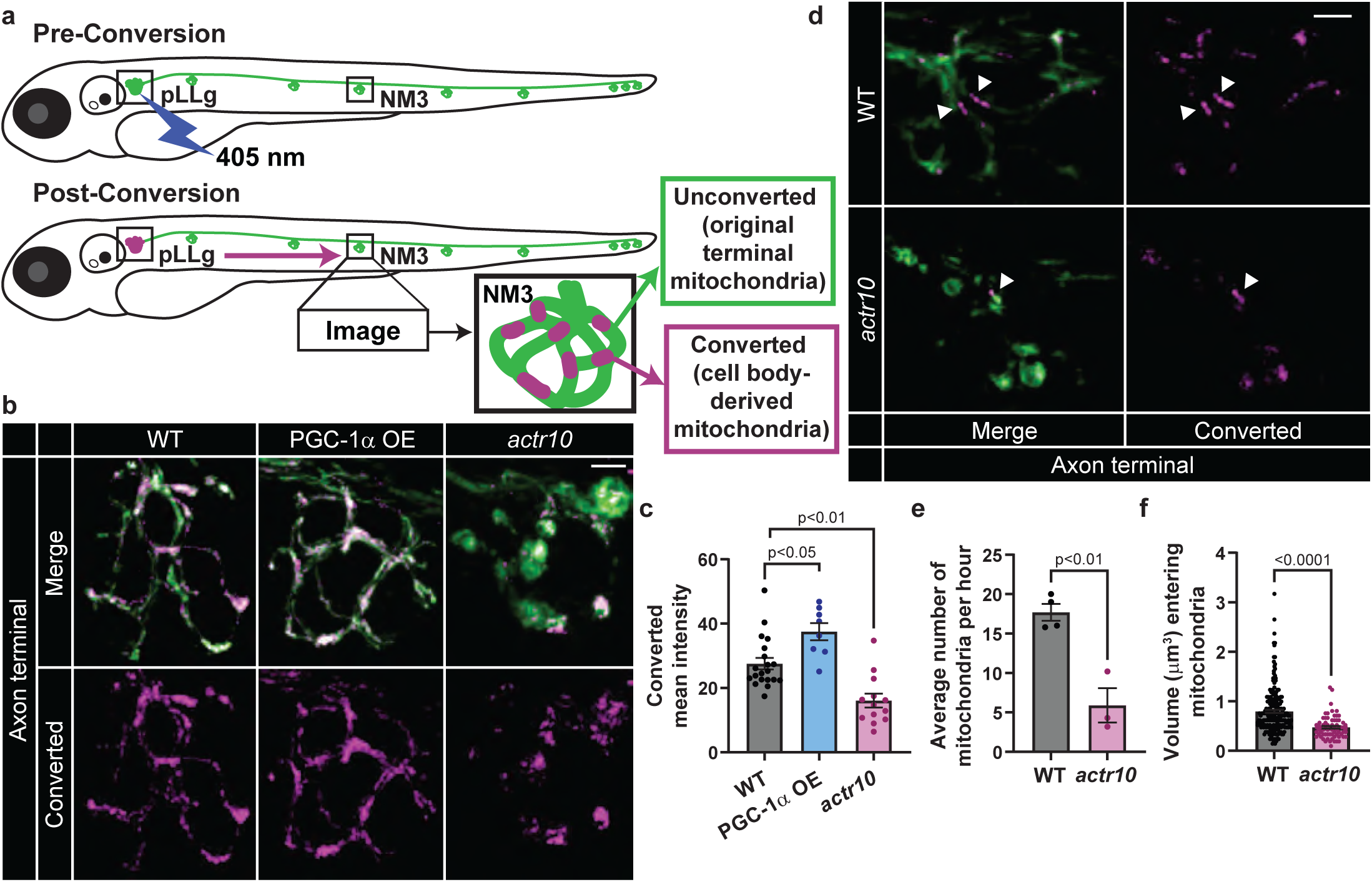
Cell body-derived mitochondrial density in the axon terminal is reduced in *actr10* mutants. (a) Photoconversion strategy. Mitochondrial matrix-localize mEos in the pLLg was locally converted using 405 nm laser. An axon terminal in the mid-trunk (neuromast 3 - NM3) was imaged 4 hpc (b) or every minute for 6h (d). (b) NM3 4 hpc in wild type, *actr10*, and transgenic overexpressing (OE) PGC-1α in neurons *(Tg(-5kbneurod1:PGC1*α*-2A-mRFP)^uwd^*^9^). (c) Mean fluorescence intensity of cell body-derived mitochondria (Converted) in the axon terminal population (Steel-Dwass). (d) Representative timepoint of a pLL axon terminal from 6 hr timelapse video (Movies S5 and S6). Cell body-derived mitochondria indicated with white arrowheads. (e, f) Quantification of the number (e, ANOVA) and size (f, Wilcoxon) of converted mitochondria entering the axon terminal. Scale bars = 5 μm. All data are mean ± SEM. Data points represent individual larvae in c and e and individual mitochondria measured from 4 wild type and 3 *actr10* axon terminals in f.

Reduced cell body derived mitochondrial material in the axon terminal could be due to either fewer mitochondria or smaller mitochondria being transported to the terminal. To differentiate this, we tracked mitochondrial addition to the axon terminal using time-lapse imaging.

Mitochondria in the pLL ganglion carrying matrix localized mEos were photoconverted and the axon terminal was imaged every minute for 6 hours (Supplementary Movies 5 and 6). *actr10* mutants had both fewer and smaller cell body derived mitochondria entering the terminal per hour (Fig. 3d-f). Together, this suggests mitochondrial biogenesis in the cell body contributes to both number and size of new mitochondria transported to the axon terminal.

### Disrupted mitochondrial fission does not alter mitochondrial biogenesis signatures

At the light microscopy level, mitochondria in *actr10* mutant pLL cell bodies appear to have a more circular and aggregated morphology, which can be indicative of impaired fission (see Fig. 1b; ^53,54^). To better understand the structure of pLL neuron cell body mitochondria, we used TEM to assess their ultrastructure in *actr10* mutants. In wildtype pLL cell bodies, mitochondria were evenly distributed throughout the cytosol (Supplementary Fig. 5a). In *actr10* mutants, cell body mitochondrial density and morphology were variable: some cell bodies had fewer but significantly enlarged mitochondria (Supplementary Fig. 5b; 5/12 cell bodies), some showed a more even distribution similar to wildtype (Supplementary Fig. 5c; 1/12 cell bodies), and some showed no visible mitochondria (Supplementary Fig. 5d; 6/12 cell bodies). In contrast, even wildtype cell bodies with little visible cytosolic area still had mitochondria present (Supplementary Fig. 5e; 0/10 cell bodies with no visible mitochondria). On average, mitochondria were significantly larger with a reduced number in *actr10* cell bodies (Supplementary Fig. 5f, g). Altered mitochondrial structure could suggest impaired mitochondrial health. However, previous work has shown *actr10* mutants have normal mitochondrial health measures in the cell body, and our TEM images show normal cristae even in the enlarged mitochondria (Supplementary Fig. 5b; ^37^). Together, these data show that *actr10* mutant mitochondria are larger and rounder than those in wild type siblings, despite not showing altered measures of health and productivity ^37^.

Altered mitochondrial fission can result in large, circular organelles ^53,54^. Given the relationship between mitochondrial fission and mitochondrial biogenesis, we asked if disrupted mitochondrial fission in *actr10* mutants could underlie disrupted mitochondrial biogenesis ^11,12^. First, we assessed length of moving and stationary mitochondria in pLL axons in *actr10* mutants and wild type siblings as a proxy for fission dynamics. These data showed no change in mitochondrial length, suggesting no major changes in fission or fusion dynamics in the axon (Supplementary Fig. 5h). The cell body mitochondria are too dense for us to directly measure their length or assess fission frequency. Instead, we disrupted mitochondrial fission and assessed measures of mitochondrial biogenesis. To disrupt mitochondrial fission, we expressed a dominant negative form of Drp1 which mimics a constitutively GDP-bound form of the enzyme (Drp1^K38A^; ^54,55^). In cultured cells, expression of Drp1^K38A^ disrupts mitochondrial fission ^54,55^.

When expressed in pLL neurons, Drp1^K38A^ causes clustering of mitochondria to the perinuclear region of the cell body and reduces mitochondrial density in both wild type and *actr10* cell bodies (Fig. 4a, b). However, Drp1^K38A^ expression does not change measures of mitochondrial biogenesis including *tfam* expression or the number of SSBP1 labeled replicating mtDNA nucleoids (Fig. 4c-f). These data argue that loss of fission is not causal in the *actr10* mutant mitochondrial biogenesis phenotype.

**Figure 4:**
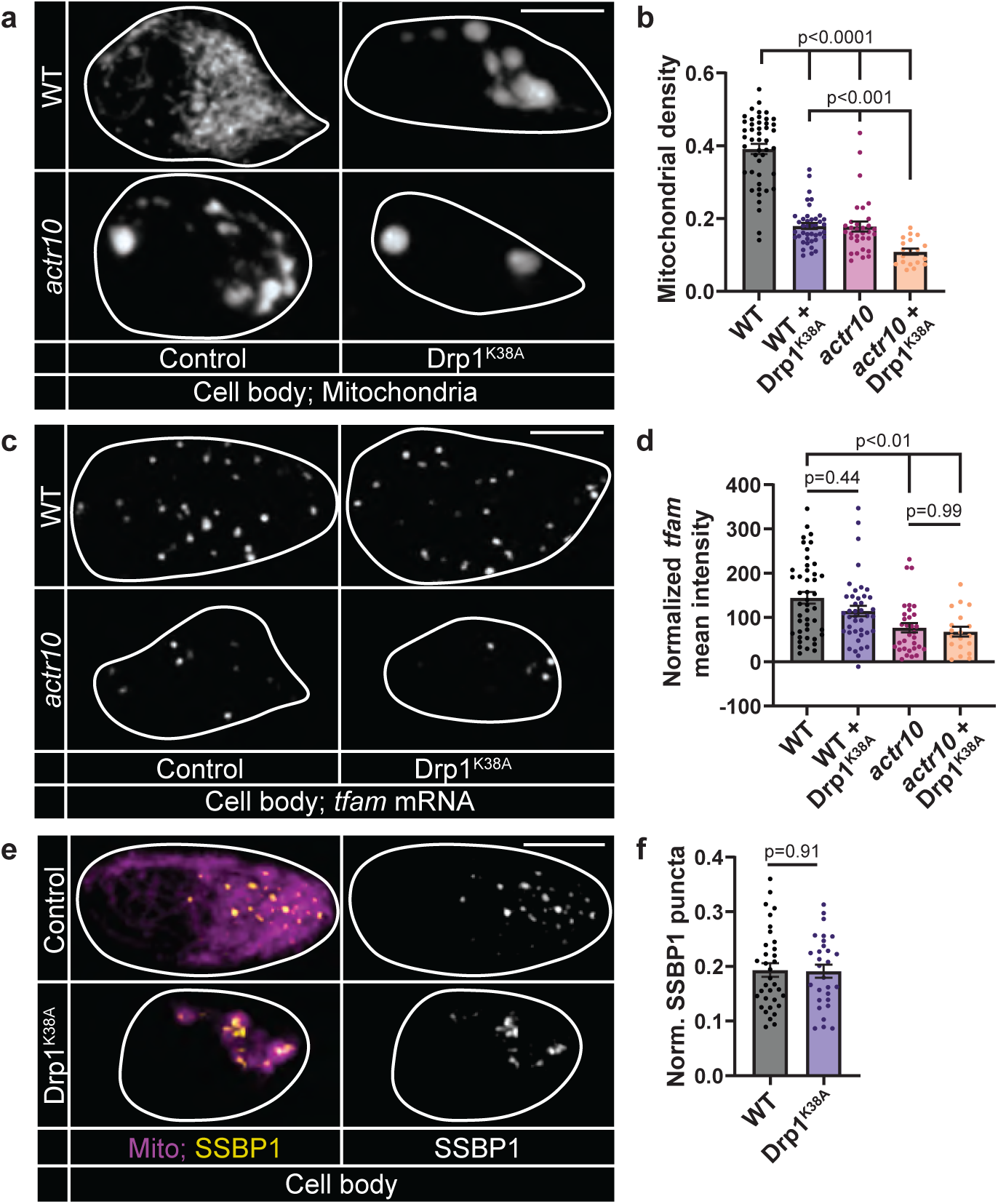
Disrupting mitochondrial fission does not alter *tfam* expression or mtDNA replication. (a) Single pLL cell bodies (outlined) expressing mitoTagRFP in wild type and *actr10* mutant larvae expressing Drp1^K38A^-RFP or RFP (control). (b) Quantification of mitochondrial density (mitochondrial area/cytosolic area; Steel-Dwass). (c) HCR RNA FISH labeling of *tfam* mRNA in a pLL cell body (outlined). (d) Mean fluorescence intensity of *tfam* normalized to background (Steel-Dwass). (e) SSBP1 immunostaining in pLL cell body mitochondria, visualized with mitochondria-localized GFP (Mito, magenta; cell outlined). (f) Number of SSBP1 puncta normalized to mitochondrial volume (ANOVA). All data are mean ± SEM. Data points represent individual larvae.

### MitoTruck mediated disruption of mitochondrial transport inhibits mitochondrial biogenesis

Our data demonstrate that inhibiting mitochondrial return to the cell body impairs mitochondrial biogenesis. To test this in another model, we engineered a synthetic tether, similar to one previously described in *C. elegans,* to constitutively move mitochondria out of the cell body and inhibit their ability to return to the cell body from the distal axon (MitoTruck; ^56^). For this, we tethered mitochondria to a constitutively active kinesin motor domain derived from KIF1A using the OMP25 outer mitochondrial membrane localization sequence (Fig. 5a; ^57,58^). When expressed in pLL neurons, MitoTruck localizes to mitochondria and ectopically concentrates these organelles to axon ends, demonstrating anterograde transport bias (Fig. 5b). Constitutive expression of MitoTruck caused failed axon extension and neuronal death by 4 dpf. To avoid any phenotypes due to failed neuronal health, we analyzed mitochondrial parameters after 24 hours of MitoTruck expression, at 2 dpf. Mitochondria in the cell body are significantly depleted in MitoTruck expressing neuron cell bodies by 2 dpf (Fig. 5c, d). This depletion is similar to that observed in *actr10* mutants at 4 dpf (mitochondrial density: WT 2 dpf 0.57 ± 0.018, MitoTruck 2 dpf 0.33 ± 0.016, WT 4 dpf 0.55 ± 0.027, *actr10* 4 dpf 0.26 ± 0.028). Analyses of mtDNA replication and *tfam* expression demonstrated that MitoTruck significantly decreased these markers of mitochondrial biogenesis (Fig. 5e-h). Together our *actr10* mutant and MitoTruck analyses demonstrate that inhibiting mitochondrial return to the cell body from the axon is sufficient to impair markers of mitochondrial biogenesis.

**Figure 5:**
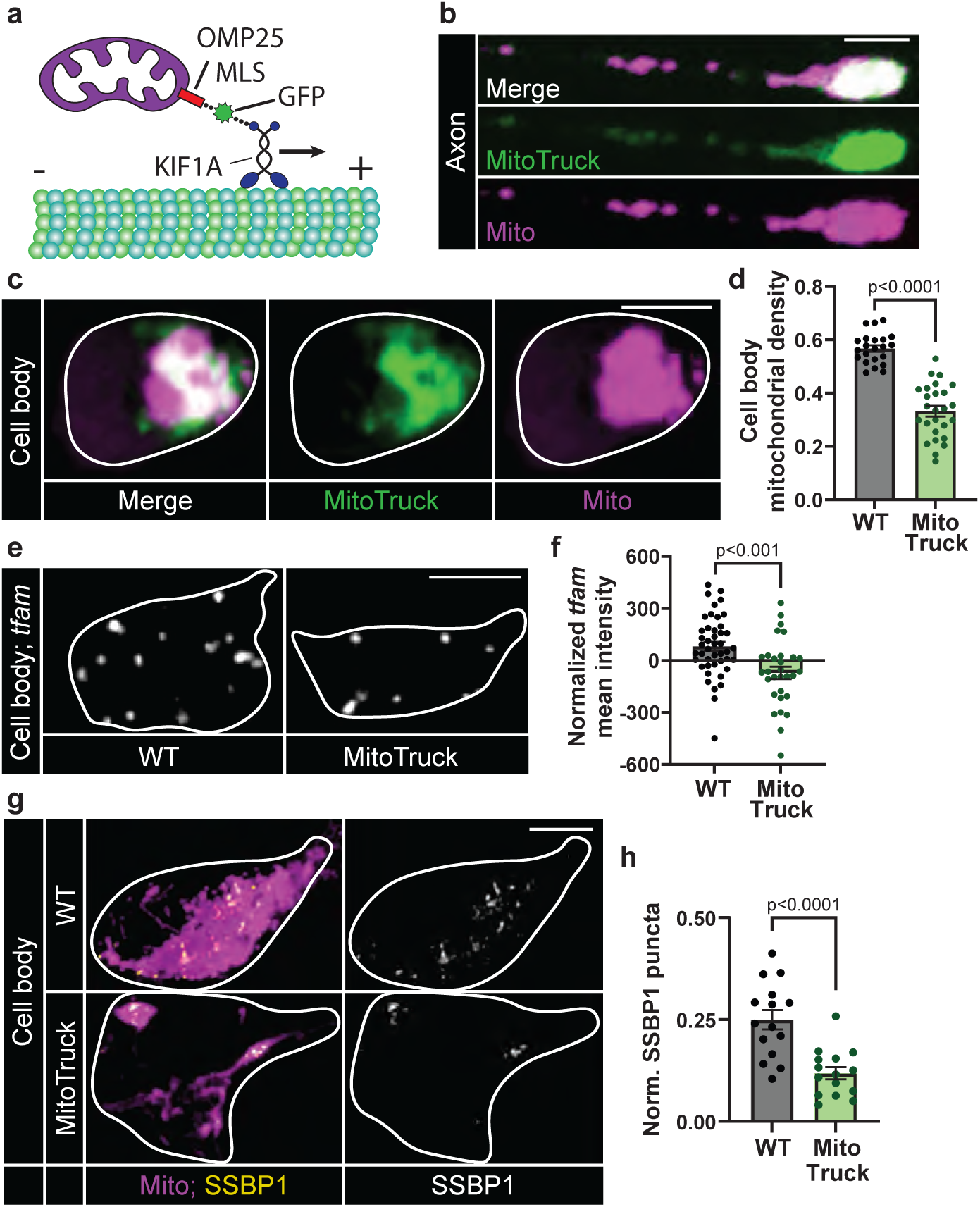
MitoTruck expression decreases cell body mitochondrial density and markers of mitochondrial biogenesis. (a) Schematic of the MitoTruck construct. The constitutively active KIF1A motor domain was tethered to mitochondria with the OMP25 outer mitochondrial membrane localization signal (MLS) and visualized with GFP. (b,c) pLL axon (b) and cell body (c; outlined) expressing MitoTruck (green) and MitoTagRFP (Mito, magenta). (d) Quantification of pLL cell body mitochondrial density (mitochondrial area/cytosolic area; Wilcoxon). (e) Representative images of HCR RNA FISH labeling of *tfam* mRNA in a single pLL cell body (outlined) without and with MitoTruck expression. (f) Mean fluorescence intensity of *tfam* normalized to background (ANOVA). (g) SSBP1 immunostaining in a single pLL cell body (outlined). Mitochondria visualized with mitochondrially localized TagRFP (Mito, magenta). (h) Number of SSBP1 puncta normalized to mitochondrial volume (ANOVA). Scale bars = 5 μm. All data are mean ± SEM and points represent individual larvae.

### Estrogen-related receptors bridge mitochondrial transport with transcriptional regulation of mitochondrial biogenesis

Our data indicate retrograde transport is required to promote mitochondrial biogenesis through regulation of upstream factors including nuclear gene expression. Several nuclear transcription factors have been identified that bind the promoters of a wide range of mitochondrial genes, allowing for coordinated expression of these genes during biogenesis ^10,59,60^. We hypothesized that a key transcription factor could link mitochondrial retrograde transport to mitochondrial biogenesis. To identify this transcription factor, we performed transcription factor enrichment analysis using significantly downregulated genes from our RNA-seq dataset. This in silico analysis identified Estrogen related receptors (ERRs) as top candidates (Fig. 6a and Supplementary Fig. 6a). ERRs have been shown to be coactivated by PGC-1α to regulate mitochondrial biogenesis ^59,61–63^. Other transcription factors regulated by PGC-1α such as NRF1/2 or PPARα were not found to be enriched in our analysis, suggesting ERRs specifically may be affected in *actr10* mutants. To validate this result, we used gene set enrichment analysis of transcription factor binding targets and identified the ERR binding motif ERR1 enriched near downregulated genes (Supplementary Fig. 6b). Finally, we took a more targeted approach by comparing our list of downregulated genes to ESRRG targets identified by ChIP-seq in cultured neurons ^63^. We found significant enrichment for known neuronal ESRRG target genes in our data set (Fig. 6b). In contrast, there was only one target gene from the Chip-seq dataset that was significantly upregulated in our dataset (p=0.885, Fisher’s exact test). Together, these in silico analyses suggest ERR-dependent transcription is reduced in *actr10* mutants.

**Figure 6:**
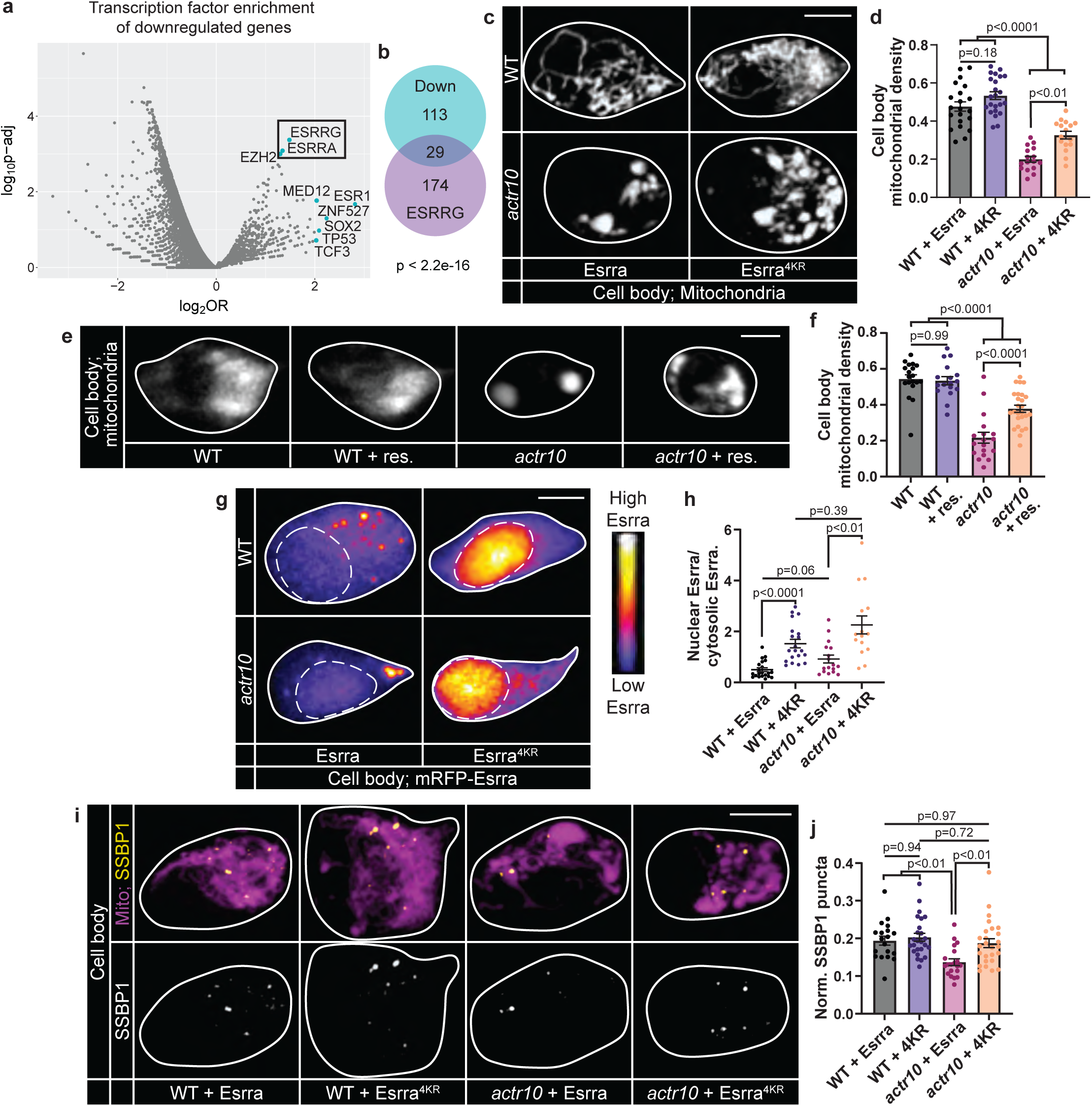
ERR activity links mitochondrial transport and mitochondrial biogenesis. (a) Transcription factor enrichment analysis for genes significantly downregulated in *actr10* mutants. Plot shows log2(Odds Ratio) vs. log10(adjusted p-value). Top 10 enriched genes based on euclidean distance from log2(OR)/log10(adj. p-value) to the origin are labeled. Box indicates ERR transcription factors identified in the dataset. (b) Overlap between downregulated genes and ESRRG neuronal ChIP-seq targets (Fisher’s exact test). Human homologs of downregulated zebrafish genes were used for transcription factor analyses. Duplicated zebrafish paralogues were only counted once in the analysis. (c) Mitochondria in wild type and *actr10* pLL cell bodies (outlined) expressing Esrra or Esrra^4KR^. (d) Quantification of mitochondrial density (ANOVAs; Tukey HSD). (e) Mitochondria in wild type and *actr10* pLL cell bodies (outlined) with and without resveratrol (+ res.) treatment. (f) Quantification of mitochondrial density (ANOVAs; Tukey HSD). (g) Heat map of mRFP-Esrra or mRFP-Esrra^4KR^ in wild type and *actr10* pLL cell bodies (outlined). (h) Mean fluorescence intensity of nuclear mRFP-Esrra/cytosolic mRFP-Esrra (Steel-Dwass). (i) SSBP1 immunostaining in pLL cell body mitochondria, visualized with mitochondria-localized GFP (Mito, magenta; cell outlined). (j) Number of SSBP1 puncta normalized to mitochondrial volume (ANOVAs; Tukey HSD). (d, f, h, j) Data points represent individual larvae; data are mean ± SEM. Scale bars = 5 μm. Mitochondrial density = mitochondrial area/cytosolic area.

If disruption of ERR-dependent transcription underlies decreased mitochondrial biogenesis in *actr10* mutant neurons, increasing ERR activity should rescue this defect. To test this, we overexpressed Esrra in pLL neurons and assessed mitochondrial density and biogenesis transcriptional programs. Similar to PGC-1α overexpression (see Supplementary Fig. 4j; ^51^), Esrra overexpression significantly increased *tfam* expression (Supplementary Fig. 6c, e). Furthermore, Esrra overexpression in *actr10* mutants rescues *tfam* expression to wild type levels (Supplementary Fig. 6d, f). However, overexpression of Esrra was unable to rescue the loss of cell body mitochondrial density in *actr10* mutants, suggesting additional levels of ERR regulation are required for optimal mitochondrial biogenesis activation (Fig. 6c, d). Activation of ERR-dependent transcription is modulated by ERR or PGC-1α posttranslational modifications (PTMs). We hypothesized PTMs are required for full Esrra activity. The most well studied modifications that activate this pathway are SIRT1-dependent deacetylation of ERR and/or PGC-1α and AMPK-dependent phosphorylation of PGC-1α ^64–68^. To determine if SIRT1 or AMPK activation were upstream of ERR-dependent mitochondrial biogenesis in neurons, we treated wild type and *actr10* mutant zebrafish with resveratrol (activator of SIRT1) or AICAR (activator of AMPK) at concentrations previously optimized for use in zebrafish larvae ^64,69,70^. Resveratrol, but not AICAR, treatment was sufficient to rescue cell body mitochondrial load in *actr10* mutant neurons (Fig. 6e, f and Supplementary Fig. 6g, h), suggesting SIRT1-mediated deacetylation of ERRs may be disrupted in *actr10* mutants.

SIRT1 has been shown to deacetylate human ESRRA at 4 conserved lysine residues (amino acids 129, 138, 160, and 162) within the DNA-binding domain. This deacetylation can be mimicked by mutating the 4 lysine residues to arginine (4KR), which conserves the basic charge of lysine but prevents acetylation^67^. Overexpression of Esrra^4KR^ in pLL neurons was able to rescue *actr10* cell body mitochondrial density (Fig. 6c, d), similar to rescue by resveratrol, suggesting deacetylation is critical for upregulation of mitochondrial biogenesis by Esrra. This was further supported by the localization of Esrra protein. Wild type Esrra did not display a canonical nuclear localization^71^ and instead remained largely cytosolic. In contrast, Esrra^4KR^ displayed strong nuclear localization, suggesting deacetylation is critical for nuclear localization to subsequently activate mitochondrial biogenesis pathways (Fig. 6g, h). Further supporting a role for deacetylation in Esrra-dependent mitochondrial biogenesis, overexpression of Esrra^4KR^ fully rescues SSBP1 punctal density in *actr10* mutants (Fig. 6i, j). These data demonstrate that ERR deacetylation is critical for regulation of mitochondrial biogenesis in pLL neurons.

SIRT1 is a nuclear-localized protein that is activated by elevated NAD+ levels^72^. To determine if NAD+ levels were altered in *actr10* nuclei, we used a genetically encoded NAD+ sensor localized to the nucleus^73^. Analysis of steady state NAD+ levels revealed a significant reduction in nuclear NAD+ in *actr10* mutants (Fig. 7a, b). A significant pool of cellular NAD+ is stored in mitochondria and changes to NAD+/NADH levels in mitochondria can affect the cytosolic/nuclear pool ^74–76^. We reasoned that altered mitochondrial transport could change local mitochondrial NAD+ levels, leading to a reduction in local SIRT1 activation in the cell body. To test this, we assessed mitochondrial NAD+ in the cell body and axon terminal of pLL neurons by localizing the genetically encoded biosensor to the mitochondria matrix. We found significantly elevated NAD+ levels in axon terminal mitochondria in *actr10* mutants while cell body mitochondria have a significant reduction in mitochondrial NAD+ levels compared to wild type controls (Fig. 7c-e). Finally, we asked if retrograde transport was responsible for moving mitochondria with higher NAD+ measures from the axon terminal to the cell body. Using live imaging, we assessed mitochondrial NAD+ levels in moving and stationary organelles using the same NAD+ sensor. These experiments demonstrated that retrogradely moving mitochondria have significantly higher NAD+ levels than either stationary or anterogradely moving mitochondria (Fig. 7f, g). Together, our data support a model in which the mitochondria with high levels of NAD+ returning to the cell body from the axon elevate local NAD+ levels to activate SIRT1-dependent ERR transcriptional activity necessary for sustained mitochondrial biogenesis in neurons (Fig. 7h). We propose that disrupting retrograde mitochondrial transport locks mitochondria with high NAD+ in the axon terminal, blocking this critical signaling step for the positive regulation of mitochondrial biogenesis.

**Figure 7:**
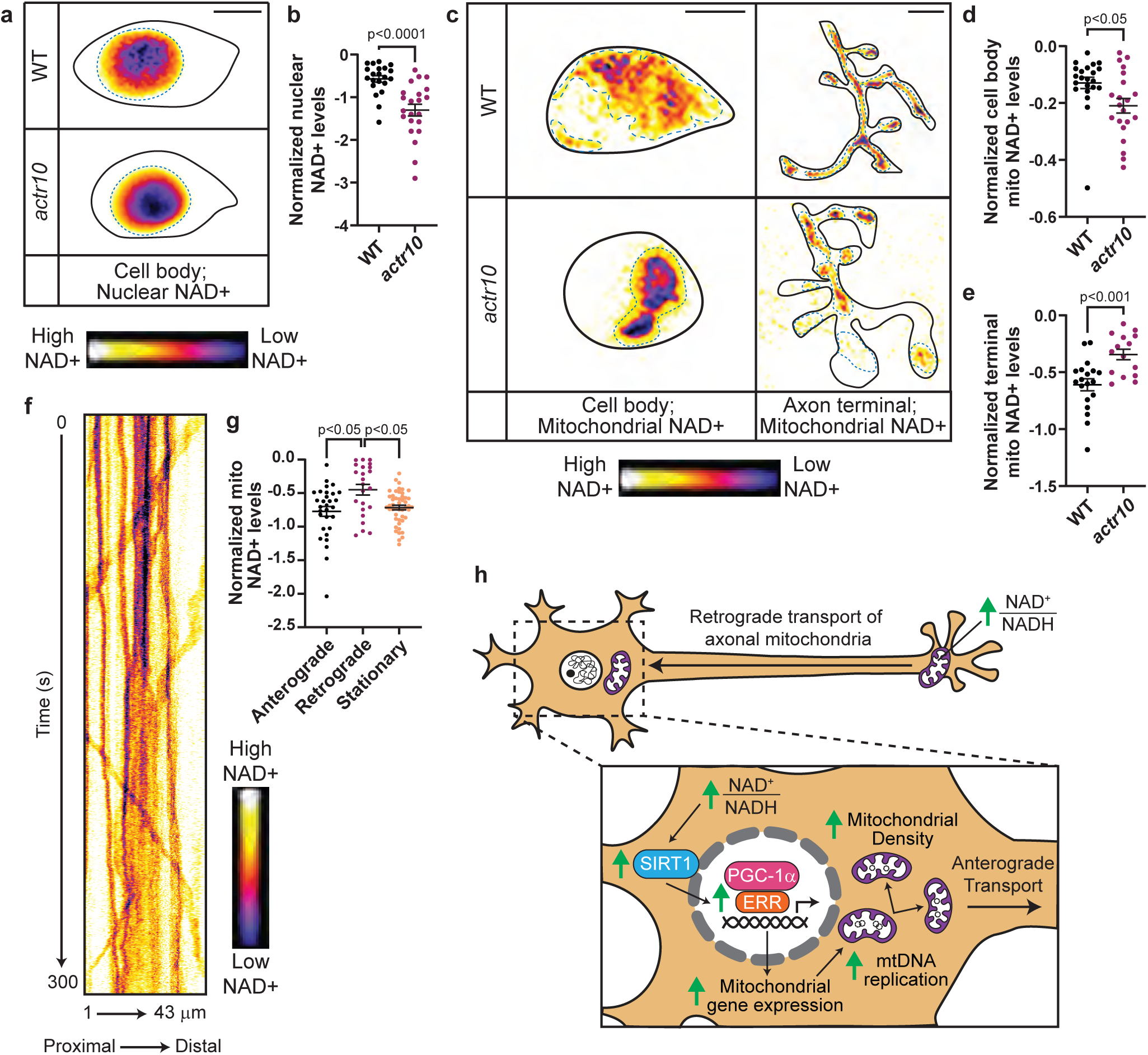
NAD+ levels are altered in *actr10* mutants. (a) pLL cell body expressing a nuclear-localized NAD+ sensor. Black outline surrounds single pLL cell body. Dotted blue outline surrounds nucleus. (c) pLL cell body and axon terminal expressing a mitochondrial matrix-localized NAD+ sensor. Black outline surrounds single pLL cell body and axon terminal. Dotted blue outline surrounds mitochondria. (f) Kymograph of pLL axon expressing a mitochondrial matrix-localized NAD+ sensor. (a, c, f) Inverted heat map indicates levels of NAD+. (b, d, e, g) Mean fluorescence intensity of NAD+ sensor normalized to cytosolic RFP (b, d, e) or mitochondrial matrix-localized TagRFP (g). Data inverted to reflect the direction of change in NAD+ levels as fluorescence intensity decreases in the presence of NAD+ (b, d: Wilcoxon; e: ANOVA; g: Steel-Dwass). (h) Model of mitochondrial retrograde transport regulation of mitochondrial biogenesis. Data are mean ± SEM. (b,d, e) Data points represent individual larvae. (g) Data points represent individual mitochondria from 14 wild type larvae. Scale bars = 5 μm.

## Discussion

Mitochondrial population maintenance is essential for neuronal health and function. These organelles are critical for a variety of functions including ATP production and calcium buffering, particularly in axon terminals ^1,2^. Through these functions, mitochondria accrue damage.

Damaged organelles must be replaced through the process of mitochondrial biogenesis. Markers of mitochondrial stress such as an increased AMP/ATP or NAD+/NADH ratio as well as increased cellular calcium can activate mitochondrial biogenesis transcriptional programs in the nucleus ^3–5,60,76–78^. However, it was unclear how these cellular signals could be transmitted long distances in the neuron to monitor mitochondrial populations in more distal regions of the neuron like the axon terminal. We identified a required role for retrograde return of axonal mitochondria to the cell body to sustain mitochondrial biogenesis transcriptional programs. We show that the return of axonal mitochondria to the cell body stimulates ERR-dependent nuclear gene transcription to regulate expression of mitochondrial genes, mtDNA replication, and ultimately increase mitochondrial biomass in the neuron. This provides a mechanism by which mitochondria in more distal regions of the neuron like the axon terminal can communicate with the nucleus to regulate mitochondrial density and mitochondrial population turnover in neurons.

The estrogen-related receptors (ESRRA, ESRRB, and ESRRG) are a family of orphan nuclear receptors first identified due to sequence similarity with estrogen receptors ^71,79,80^. The three ERR proteins bind the ERR response element (ERRE) which is found in the promoter of many mitochondrial genes ^59,71^. Because of this, increased ERR activity is associated with increased mitochondrial biogenesis in a number of tissues, including neurons ^61,62,73,81^. Despite their similarity to estrogen receptors, ERRs do not have the capacity to bind estrogen, but rather are thought to be constitutively active ^82^. Their activity can be regulated through expression level, PTMs, or interaction with co-activators like PGC-1α ^8,9,81^. PGC-1α regulates ERR transcription and also interacts with ERRs as a coactivator to induce the transcription of mitochondrial biogenesis genes ^8,9,62^.

ERRs serve as mediators between mitochondrial signals and biogenesis transcriptional programs. In non-neuronal cells, signs of mitochondrial depletion or failed health like reduced cellular energy or increased cytosolic calcium have been shown to activate ERR-dependent transcription, either directly or through PGC-1α activation ^13,83^. These upstream signals of cellular and mitochondrial health act through enzymes that can directly modify ERR and PGC-1α via PTMs including phosphorylation and acetylation. Two of the most well studied upstream regulatory factors are AMPK and SIRT1, which serve as energy sensors for the cell ^6^. AMPK is activated by an increased AMP to ATP ratios and can activate mitochondrial biogenesis through PGC-1α phosphorylation ^4,84^. SIRT1 is activated by an increased NAD+ to NADH ratio and can activate mitochondrial biogenesis through PGC-1α and/or ERR de-acetylation ^3,67^. AMPK and SIRT1 can be artificially activated through AICAR and resveratrol treatment, respectively, to induce mitochondrial biogenesis ^64,65,69,84,85^. In our experiments, resveratrol, but not AICAR, treatment was able to rescue *actr10* biogenesis defects, suggesting SIRT1-dependent mitochondrial biogenesis is lost. SIRT1 has been shown to deacetylate human ESRRA at four lysine residues to increase ERRA transcriptional activity^67^. Overexpression of an Esrra mutant mimicking deacetylation at these sites was able to rescue markers of mitochondrial biogenesis in *actr10* mutants, including mitochondrial density and mtDNA replication. Together, this suggests retrograde transport of axonal mitochondria may stimulate ERR-dependent mitochondrial biogenesis through activation of SIRT1, though we cannot rule out other upstream modulators which could function with SIRT1 in this process.

SIRT1 requires NAD+ as a co-substrate for the removal of acetyl groups from lysines, making NAD+ levels a direct regulator of SIRT1 enzymatic activity ^72^. Increased NAD+ levels have been shown to increase SIRT1-mediated de-acetylation of PGC-1α, increasing its transcriptional activity ^77,85^. Therefore, local NAD+ levels are a likely signaling mechanism by which non-neuronal cells are able to sense energy levels and respond with transcriptional regulation of mitochondrial biogenesis. However, it was unclear how the predominantly nuclear-localized SIRT1 could sense and respond to levels of NAD+ in mitochondrial populations spread throughout the expansive dendritic and axonal arbors of neurons ^86^. Our data suggests that retrograde transport is required to move mitochondria with high levels of NAD+ to the cell body to stimulate biogenesis. We show retrogradely transported mitochondria have higher NAD+ levels and that the NAD+ levels in axon terminal mitochondria are increased when retrograde transport of mitochondria is impeded. We find that failure to return mitochondria with elevated NAD+ levels to the cell body in *actr10* mutants leads to reduced nuclear and cell body mitochondrial NAD+ levels. Based on this, we propose that local SIRT1 activity in the cell body requires increases in the local NAD+ levels which is accomplished by the movement of mitochondria with elevated NAD+ from the axon back to the cell body. Consequent activation of SIRT1 in the cell body then promotes mitochondrial biogenesis through de-acetylation of transcriptional regulators like PGC-1α or ERRs. In support of this, treatment with resveratrol, which circumvents the requirement for NAD+, can rescue mitochondrial density in *actr10* mutant cell bodies. It is likely that additional energy sensing pathways work with NAD+/SIRT1 to control mitochondrial biogenesis through additional transcription factor PTMs. Further work on transcription factor PTMs and associated modulatory pathways will clarify the role of energy sensing through NAD+ and other signaling molecules on the control of mitochondrial biogenesis in neurons.

In conclusion, our work demonstrates that transport of mitochondria from the axon back to the cell body is a positive regulator of neuronal mitochondrial biogenesis through regulation of ERR-dependent gene transcription. Mitochondria derived from biogenesis in the neuronal cell body are critical for maintaining cell body mitochondrial density as well as supplying the anterogradely transported mitochondria that support the axon terminal mitochondrial population. Unsurprisingly, impaired mitochondrial biogenesis can diminish neuronal health and has been linked to diseases like Parkinson’s and Alzheimer’s diseases ^87–90^. However, regulatory mechanisms that govern this critical process in the neuron remain poorly understood. Here we show that neurons in zebrafish larvae require mitochondrial retrograde transport for effective mitochondria-nuclear communication to regulate mitochondrial biogenesis through ERRs. This mechanism provides new insight into how neurons regulate mitochondrial population health and density in response to changing demands on the organelle in distal compartments.

## Acknowledgments

We would like to acknowledge members of the Drerup lab and Dr. Samantha Lewis for thoughtful discussions of the work and critique of the manuscript. The constitutively active KIF1A motor domain construct was provided by Dr. Marvin Bentley. We would like to thank the University of Wisconsin Carbone Cancer Center Flow Cytometry Laboratory, supported by P30 CA014520, for use of its facilities and assistance with fluorescence activated cell sorting. We would like to thank Randall Massey and the University of Wisconsin School of Medicine and Public Health Electron Microscopy Facility for assistance with Transmission Electron Microscopy.

## Author Contributions

Conceptualization: AEL, CMD

Data acquisition and analysis: AEL, CS, RS, CMD

Funding acquisition: AEL, CMD

Writing – original draft: AEL

Writing – review & editing: CMD

## Data and materials availability

All data and materials are available upon request from the corresponding author.

## Materials and Methods

### Zebrafish husbandry

All zebrafish (*Danio rerio*) work was done in accordance with the University of Wisconsin-Madison Institutional Animal Care and Use Committee guidelines. Adult zebrafish were kept at 28°C and embryos were spawned according to established protocols ^91^. Embryos and larvae were kept in embryo media (995 μM MgSO4, 154 μM KH2PO4, 42 μM Na2HPO4, pH 7.2, 1.3 mM CaCl2, 503 μM KCl, 15 mM NaCl, 714 μM NaHCO3), maintained at 28°C, and developmentally staged using established methods ^92^. All experiments were performed at 4 dpf unless otherwise specified. Sex is not determined at this stage. Zebrafish strains used for these experiments are listed in Table 1. The Tg(*hsp70l:EGFP-en.sill*)*^uwd^*^12^ transgenic was derived using Tol2-mediated transgenesis and the *hsp70:eGFP-SILL* plasmid as described ^93^. Genotyping for *actr10^nl^*^15^ ^35^, *p150a^y^*^625^ ^37^, *p150b^nl^*^16^ ^35^, and *nudc^nl^*^21^ ^42^ was done as previously described using the primers in Table 2.

### Cloning

The DNA expression plasmids used in this paper are listed in Table 1. Novel DNA expression constructs were constructed using Gateway cloning^93^ or Gibson cloning with oligonucleotides listed in Table 2 ^94^. *5kbneurod:POLG2-GFP-p2a-mitoTagRFP* was created from a human POLG2-GFP template plasmid^12,46^. *5kbneurod1:kif1a-egfp-omp25* [“MitoTruck”] was created using the zebrafish outer membrane protein 25 (OMP25) mitochondrial localization sequence (MLS) cloned from 4 dpf zebrafish cDNA. The truncated constitutively active rat ortholog of KIF1A was cloned from *pBA-kif1a393-tdTomato-FKBP* ^95^. *5kbneurod1:cox8a-cox8a-halotag* was created by fusing two copies of the cox8a MLS to a HaloTag. *5kbneurod1:esrra-p2a-mRFP* was created using the zebrafish *esrra* sequence derived from 4 dpf zebrafish cDNA and existing p2a and mRFP plasmids^93^. Esrra^4KR^ expression plasmid was created using consecutive site-directed mutagenesis reactions with the Agilent QuikChange Lightning Site-Directed Mutagenesis Kit. Lysines at amino acid 129, 138, 160, and 162 in the zebrafish ortholog of Esrra were converted to arginines. *5kbneurod1:cox8a-cox8a-NAD-p2a-RFP* was created from plasmids containing a double cox8a MLS, a NAD+ biosensor (Addgene plasmid 186791; ^96^) and a p2a-RFP. *5kbneurod1:H2B-NAD-p2a-RFP* and *5kbneurod1:cox8a-cox8a-NAD-p2a-mitoTagRFP* were created from the *5kbneurod1:cox8a-cox8a-NAD-p2a-RFP* plasmid and either a plasmid containing an H2B nuclear localization signal or a p2a-mitoTagRFP.The NAD+ biosensor consists of a circularly permuted Venus fluorescent protein (cpVenus) with a bipartite NAD+-binding domain modeled from bacterial DNA ligase ^96^. *5kbneurod1:cox8a-cox8a-paGFP-p2a-mitoTagRFP* was created from plasmids containing a double cox8a MLS, a photo-activatable GFP, and a p2a-mitoTagRFP. *hsp70l:EGFP-en.sill* was created using existing plasmids containing the hsp70 minimal promotor, eGFP, and the 3’ SILL enhancer element that limits expression to the pLL ganglion ^93,97^. *hsp70l:POLG2-GFP-p2a-mitoTagRFP* was created from the *5kbneurod:POLG2-GFP-p2a-mitoTagRFP* plasmid and the hsp70 minimal promoter. For all expression plasmids except the *hsp70* driven constructs, expression was controlled by a 5kb portion of the *neurod* promotor as previously described ^93,98^. All novel DNA expression plasmids were sequenced and verified prior to use.

### Live imaging acquisition and analysis

For live imaging of pLL neurons, we used a combination of transient transgenesis and stable transgenic expression. For transient transgenesis, 3-25 pg of plasmid DNA was microinjected into zebrafish zygotes as previously described ^35,99,100^. Plasmids used for these experiments are listed in Table 1. For imaging, zebrafish larvae were sorted to identify larvae with expression in a subset of pLL neuron cell bodies using an AxioZoom V.16 Zeiss microscope. For HaloTag constructs, larvae were treated with Janelia Fluor 635 in embryo media at 1 μM for 1-2 h or 0.1 μM overnight at 28.5°C in the dark. For all live imaging, larvae were anesthetized in 0.02% tricaine, mounted in 1.5% low melt agarose in embryo media, and imaged with an Olympus FV3000 confocal microscope with a 40x (NA 1.25) silicone oil objective. Optimal interslice interval was used for all imaging.

Larval TgBAC(*neurod:egfp*)*^nl^*^1^ (Fig. 1a) and *Tg(hsp70:eGFP-SILL*)*^uwd12Tg^* (Supplementary Fig. 1a) transgenic zebrafish images were created by stitching overlapping regions of the larvae imaged using the Olympus FV3000 with a 10X objective using the pairwise stitching plugin in ImageJ ^101,102^. Non-linear adjustment to midtones to show the full nervous system was done using the levels function in Adobe Photoshop. For TgBAC(*neurod:egfp*)*^nl^*^1^, a brightfield image of the full larva was taken using the AxioZoom V.16 Zeiss microscope with a Zeiss Axio 506 color camera and overlaid using Adobe Photoshop with opacity adjusted to 10%. For *Tg(hsp70:eGFP-SILL*)*^uwd12Tg^*, brightfield images from the Olympus FV3000 were stitched and overlaid with an opacity of 30%.

To quantify mitochondrial density in the pLL cell body and axon terminal, area was measured from standard deviation z-projections in ImageJ. Mitochondrial area was measured using a manually applied threshold while cytosolic area was measured by manual tracing or applied threshold of a cytosolic fill. Mitochondrial density was calculated as mitochondrial area divided by cytosolic area. To quantify mitochondrial or cell volume, the 3D objects counter was used in ImageJ using a manually applied threshold ^102^.

For POLG2 axon terminal analysis, *5kbneurod:POLG2-GFP-p2a-mitoTagRFP* was transiently expressed in pLL neurons by zygotic microinjection as described ^100^ and terminal regions were imaged as described above. POLG2+ puncta were manually counted and mitochondrial volume was quantified as described above.

Mitochondrial axonal transport analyses were done as described previously ^100^. Briefly, transient transgenesis of *5kbneurod1:mitotagRFP* was used to visualize mitochondria in pLL axons. 30–100 μm lengths of axon were imaged as a single *z*-plane at a rate of 300 ms per frame for 1000 frames. Kymographs were generated using the ImageJ Multi Kymograph function ^102^.

Anterograde, retrograde, and bidirectional mitochondria number were quantified using kymographs while stationary mitochondria number was quantified by visual analysis of transport videos. Mitochondrial length was manually measured from transport movies using the line selection tool in ImageJ.

For NAD+ quantification, NAD sensor constructs were transiently expressed in pLL neurons by zygotic microinjection. For single timepoint imaging, larvae were mounted live as described above and cell body and axon terminal regions were imaged at 4 dpf with consistent laser power and detector gain for all images. Transport analysis was done as described above with sequential imaging of cpVenus and RFP channels^35,100^. For cell body and axon terminal mitochondria analysis (*5kbneurod1:cox8a-cox8a-NAD-p2a-RFP*), cell body and axon terminal regions were isolated using the mRFP cytosolic signal as a mask and the image subtraction function in ImageJ ^102^. cpVenus and mRFP mean fluorescence intensity were then quantified from sum projections of the isolated region. For nuclear analysis (*5kbneurod1:H2B-NAD-p2a-RFP*), sum projections were created for cpVenus and mRFP. Nuclear cpVenus mean fluorescence intensity was quantified by manual outlining of nucelar area. mRFP mean fluorescence intensity was quantified by manual outlining of cytosolic area. For transport analysis (*5kbneurod1:cox8a-cox8a-NAD-p2a-mitoTagRFP)*, mitochondria were identified as moving anterogradely, retrogradely, or stationary by visual analysis of transport videos. cpVenus and RFP mean fluorescence intensity were quantified by manual outlining of individual mitochondria. For all analyses, mean fluorescence intensity of cpVenus was normalized to mean fluorescence intensity of RFP to control for differences in expression level. NAD+ binding to the bipartite NAD+-binding domain in the biosensor decreases cpVenus fluorescence intensity in a dose-dependent manner ^96^. The images in Figure 7 were generated using the fire lookup table in ImageJ and the images/data are inverted to appropriately reflect the direction of change in NAD+ levels.

### Drug treatment

Resveratrol and AICAR (5-aminoimidazole-4-carboxamide ribonucleotide) stocks (200 mM and 27.5 mM respectively) were diluted in embryo media to concentrations previously shown to be effective in larval zebrafish (Resveratrol 300 μM, AICAR 1 mM; ^70^). At 3 dpf, transient transgenic larval zebrafish expressing cytosolic GFP and mitochondrially localized tagRFP (see Table 1 for DNA constructs) were placed in a 6 well plate and incubated overnight (18-24 h) in drug or equal amount of drug solvent (DMSO for resveratrol, sterile H_2_O for AICAR) in embryo media. After treatment, larvae were mounted live, pLL neuronal cell bodies were imaged, and mitochondrial density analyzed as described above.

### Heat shock-induced expression of Drp1^K38A^ and POLG2-GFP

For DRP-1 analysis, *Hsp701:Drp1K38A-mRFP* ^35^ or *5kbneurod1:RFP* ^35^ (control) was transiently expressed with *5kbneurod1:mito-eGFP* in pLL neurons by zygotic microinjection as described ^100^. At 4 dpf, larvae of both groups were placed in PCR strip tubes (2-3 larvae per tube) with approximately 200 μl of embryo media. Larvae were incubated at 37°C for 1 hour in a PCR machine (BioRad) then returned to 28°C. Larvae were fixed in 4% paraformaldehyde 7 hours post-heat shock for subsequent HCR RNA FISH or immunolabeling as described below.

For POLG2-GFP cell body puncta analysis, *Hsp701:POLG2-GFP-p2a-mitoTagRFP* was transiently expressed in pLL neurons by zygotic microinjection as described ^100^. At 4 dpf, embryo media was pre-heated to 37°C in 50 mL conical tubes. Larvae were transferred to the pre-heated embryo media in 50 mL conical tubes and kept at 37°C for 1 hour. Larvae were then returned to 28°C media. Larvae were fixed in 4% PFA/0.25% Triton X-100 7 hours post-heat shock for subsequent imaging. Fixed larvae were mounted between glass slides and #1.5 coverslip with Fluoromount mounting medium prior to imaging with an Olympus FV3000 confocal microscope with a 60x (NA1.42) oil objective. Punctal quantification was done as described above for axon terminals.

### Nuclear localization analysis

*5kbneurod1:mRFP-esrra* or *5kbneurod1:mRFP-esrra^4KR^*was transiently expressed in pLL neurons by zygotic microinjection as described ^100^. At 4 dpf, larvae were fixed in 4% PFA/0.25% Triton X-100 and co-incubated with DAPI (1:500) at 4°C overnight. Larvae were imaged as described above for cell body POLG2-GFP. DAPI was used to mask out nuclear RFP signal. Mean fluorescence intensity was then quantified from sum projections of the nuclear and cytosolic regions.

### Photolabeling

*5kbneurod1:cox8a-cox8a-paGFP-p2a-mitoTagRFP* was transiently expressed in pLL neurons by zygotic microinjection as described ^100^. Cell body or axon terminal mitochondrial-matrix localized PA-GFP was specifically photoactivated by illuminating with a 405 nm laser using a region of interest (ROI) defined around the structure. For each larva, a pre-activation image was taken (488 nm and 561 nm lasers), the region was scanned for 1 to 5 seconds with 1% 405 nm laser power using the stimulation function for conversion, and a post-activation image (488 nm and 561 nm lasers) was taken to confirm activation. The axon terminal was then re-imaged at set timepoints post-conversion. Due to transient expression, PA-GFP was not expressed in every axon terminal. To reduce variability due to differences in axon length, only axon terminals innervating sensory organs in the center of the trunk (neuromasts 2, 3, or 4) were used for analysis. PA-GFP and MitoTagRFP mitochondrial area were quantified as described earlier. “Proportion original mitochondria” for terminal activation and “proportion of cell body-derived mitochondria” for cell body activation was quantified as PA-GFP area/MitoTagRFP area (see Figure S3).

Photoconversion of mitochondrially localized mEos was performed using the stable transgenic *Tg(-5kbneurod1:mito-mEos)^y^*^586^*^Tg^* as previously described ^37^. The pLL ganglion was specifically converted by illuminating with a 405 nm laser through a z-stack of the entire ganglion with regional restriction using a region of interest (ROI) defined around this structure. For each larva, a preconversion image was taken (488 nm and 561 nm lasers), the stack was scanned with 1% 405 nm laser power for conversion, and a postconversion image (488 nm and 561 nm lasers) was taken to confirm complete green to red conversion of mEos in pLL cell bodies. For 4 hour timepoint analysis, the axon terminal of a sensory organ in the center of the trunk (NM3) was imaged 4 hours post pLL ganglion conversion. All microscope settings were kept consistent for all images. For axon terminal analysis, the red and green channels were merged and a mask was created by thresholding a standard deviation projection of the axon terminal *z*-stack. The mask was applied to a sum projection of the red channel and mean fluorescence was quantified. For timelapse analysis, a z-stack through an ROI surrounding NM3 was scanned minute for 6 hours. Image analysis was performed by manually scanning through standard deviation z-stack projections from the time series. Cell body derived (red photoconverted) mitochondria were counted and volume measured using the 3D objects counter in ImageJ.

### In situ Hybridization and Hybridization Chain Reaction RNA Fluorescent In situ Hybridization (HCR RNA FISH)

Colorimetric in situ hybridization was done according to established protocols ^99,103^. To generate probes for *in situ* hybridization, we amplified 200-600 base pair regions of relevant cDNA within the open reading frames of genes of interest to use as a template for probe synthesis (^104^; Table 2). Zebrafish were fixed overnight in 4% paraformaldehyde (PFA) in phosphate-buffered saline (PBS) at 4°C. Fixed larvae were washed in PBS/0.1% Tween-20 and dehydrated in a methanol series. Larvae were stored in methanol at -20°C. Larvae were rehydrated with PBS/0.1% Tween-20 in a reverse methanol series. Larvae were treated with 10 μg/ml proteinase K in PBS/0.1% Tween-20 for 15 minutes then immediately fixed for 20 min in 4% PFA at room temperature (RT). Larvae were washed with PBS/0.1% Tween-20 then prehybridized for 1-2 hours at 65°C. Larvae were incubated overnight at 65°C with probe diluted 1:200 in hybridization buffer (10 mM citric acid, 0.1% Tween-20, 50 μg/ml heparin, 5× saline sodium citrate (SSC), 50% formamide). Larvae were washed at 65°C with a hybridization buffer-SSC buffer series (2:1 buffer:2x SSC 15 min; 1:2 buffer:2x SSC 15 min; 2x SSC 15 min; 0.2x SSC 30 min twice).

Larvae were then washed at RT with 2:1 and then 1:2 0.2x SSC:PBS/0.1% Tween-20 for 5 min. Larvae were blocked in block solution (2% goat serum, 10% bovine serum albumin in PBS/0.1% Tween-20) for 2 hours at RT, and then sheep anti-digoxigenin (1:10,000) in block solution overnight at 4°C. Larvae were washed 6 times 15 min in PBS/0.1% Tween-20, 3 x 5 min in coloration buffer (100 mM Tris–HCl pH 9.5, 50 mM MgCl2, 100 mM NaCl, 0.1% Tween-20), then stained with 0.45% nitro blue tetrazolium, 0.35% 5-bromo-4-chloro-3-indolyl phosphate in a coloration buffer in dark until color development. Larvae were washed with PBS/0.1% Tween-20, fixed for 20 min in 4% PFA at RT, dehydrated in a methanol series, and rehydrated for imaging and sorting of RNA expression. WT and *actr10* mutant zebrafish were kept in the same tube for processing, then blindly sorted based on RNA expression into two groups (high and low expressing). Representative larvae were mounted in 1.5% low melt agarose in embryo media and imaged with a Zeiss Axio Imager.M2 microscope with Zeiss Axio 506 color camera. Sorted larvae were genotyped using the oligonucleotides in Table 2.

HCR RNA FISH was performed as described previously ^51^. An mRNA transcript-specific probe set for each gene of interest was designed by Molecular Instruments (Table 3). Larvae fixation, methanol dehydration/rehydration, and Pro K treatment were performed as described for colorimetric in situ hybridization with modification in the Pro K treatment for larvae less than 4 dpf. 2 dpf larvae were treated with Pro K for 5 min, 3 dpf for 10 min, and 4 dpf for 15 min, followed by post-fixation. Larvae were incubated with a manufacturer-supplied hybridization buffer at 37°C for 1 h and then incubated in 4 nM (*pgc1a* and *tfam*) or 16 nM (others) probe in hybridization buffer at 37°C overnight. Larvae were washed with a manufacturer-supplied wash buffer and then SSC/0.1% Tween-20. Then, larvae were incubated in a manufacturer-supplied amplification buffer at RT for 30 min, during which manufacture-supplied reporter-labeled fluorescent hairpins were individually heated at 95°C and then cooled to RT for 30 min in the dark. Larvae were placed in fresh amplification buffer with 60 nM hairpins. B1 probes were labeled with 546 nm wavelength hairpins while B2 probes were labeled with 647 nm wavelength hairpins combined in the same tube. Larvae were incubated overnight in the dark at RT. Larvae were mounted between glass slides and #1.5 coverslip with Fluoromount mounting medium prior to imaging with an Olympus FV3000 confocal microscope with a 60x (NA1.42) oil objective.

### For each probe, laser power and detector gain were kept consistent for all images

To quantify HCR signal in the whole pLL ganglion in *TgBAC(neurod:eGFP)*^nl1^ larvae, a mask was created using the GFP channel threshold using the default auto thresholding algorithm in ImageJ and the image subtraction function was used to remove non-neuronal signal. After using the mask to exclude signal outside the pLLg, a sum projection of the probe signal was created and mean fluorescence intensity was measured. To quantify HCR signal in single cell bodies, a substack was created around the cell of interest and a sum intensity z-projection of the probe signal was created. The cell body was manually outlined and mean fluorescence intensity of the probe signal within the cell body was measured. To quantify background signal, a region outside the pLLg or cell body was selected and the sum intensity of probe signal was measured. Background mean signal was subtracted from pLL mean signal to get the normalized mean fluorescence intensity.

### Immunolabeling

For SSBP1 and TFAM immunolabeling, transient transgenesis of either *5kbneurod1:mito-eGFP* or *5kbneurod1:mitotagRFP* was used to localize pLL cell body mitochondria. Larvae were fixed in 4% PFA/0.25% Triton X-100 at 4°C overnight then incubated in water at RT overnight. Larvae were kept in block (0.1% Triton X-100, 1% dimethyl sulfoxide, 0.02% sodium azide, 0.5% bovine serum albumin, 5% goat serum) at RT for 2h, then incubated with antibody in block overnight at 4°C. Antibodies were validated by western blot (Supplementary Fig. 1m). Larvae injected with *5kbneurod1:mito-eGFP* were incubated with chicken anti-GFP (1:2000) to amplify signal. Larvae were incubated with either rabbit anti-SSBP1 (1:500) or rabbit anti-TFAM (1:100). Larvae were then washed in PBS/0.1% Triton X-100 before incubation in goat anti-chicken AlexaFluor 488 (1:1000) and goat anti-rabbit AlexaFluor 568 (1:1000) or goat anti-rabbit AlexaFluor 647 (1:1000) at 4°C overnight. Larvae were washed in PBS/0.1% Triton X-100 prior to imaging as described for HCR. For analysis, a mask was created from the mitochondrial channel using the default auto thresholding algorithm in ImageJ and the image subtraction function was used to remove signal outside the pLL cell body. For TFAM quantification, the 3D objects counter function in ImageJ and manual thresholding was used to quantify volume of TFAM puncta and mitochondria (mitochondrial volume used for normalization). For SSBP1 quantification, SSBP1 puncta were manually counted and normalized to mitochondrial volume.

### Transmission Electron Microscopy

4 dpf wild type and *actr10* mutant larvae samples were dissected by removing the trunks posterior to the pLLg and then fixed in Karnovsky’s fixative (2.5% glutaraldehyde/2.0% formaldehyde in 0.1M NaPO4 buffer (PB), pH=7.2) at 4°C overnight. All remaining steps were performed on the specimens in glass scintillation vials on a rotator unless specified. The fixative was removed and the samples washed 5 x 5 min in PB at RT. Post-fixation was performed on the samples in 1% OsO4 in PB for 1 hour at RT. The samples were again washed in PB, 7 x 5 minutes at RT. Dehydration was performed with a graded series of EtOH (35%, 50%, 70%, 80%, 90% 5 x 5 minutes, 95% for 10 minutes, 100% for 3 x 10 minutes) at RT. Specimens were transitioned from EtOH by rinsing 2 x 7 minutes in acetone at RT. Samples were infiltrated and embedded in EMBed 812 resin (EMS Hatfield; PA 19440) using a graded series of 2:1 Acetone /

EMBed812 1 hour RT, 1:1 Acetone / EMBed812 overnight RT, 1:2 Acetone / EMBed812 2 hours RT. 100% EMBed 812 resin infiltrations continued in uncapped vials for 1 hour at 65°C in a laboratory microwave, with a fresh change of resin for 1 hour at 75°C. Final embedding was done in EMBed 812 resin in aluminum weighing dishes placed in a 60°C drying oven and allowed to polymerize for 2 days. Polymerized samples had the aluminum dishes stripped, with the target specimen area sawed out with a jewelers saw and glued to blank polymerized stubs of EMBed 812 resin. Specimens were sectioned on a Reichert-Jung Ultracut E ultra-microtome with sections collected on formvar coated, Cu 2×1 slot grids. Grids were post-stained in uranyl acetate and Reynolds lead citrate and viewed on a Philips CM120 at 80kV. Images were collected on a BioSprint12 digital camera. One section of the pLLg was imaged for a WT and *actr10* mutant larvae. All cells visible within the section were analyzed. Images were calibrated in ImageJ and analyzed by manual counting of mitochondria and manual outlining for mitochondrial, cell, and nuclear area. Cytosolic area was calculated as cell area minus nuclear area.

### Western blot

Western blot of TFAM and SSBP1 protein was done as described previously ^35^. Protein lysates of 4 dpf wildtype larvae were run on a 12% sodium dodecyl sulfate polyacrylamide electrophoresis gel. After transfer to the polyvinylidene difluoride membrane, the blots were blocked in 5% nonfat dry milk in 1X PBS/0.1% Tween for 1–4 hours before incubation with anti-TFAM (1:1000) or anti-SSBP1 (1:500) overnight in block at 4°C. Membranes were then washed in 1X PBS, 0.1% Tween20 and incubated with secondary antibody conjugated to horseradish peroxidase for 90 min at RT. The blot was developed with a Western Sure Chemiluminescent Substrate (LiCor) on a C-DiGit Blot Scanner.

### Embryo dissociation and fluorescence activated cell sorting (FACS)

20-30 4 dpf *Tg(hsp70:eGFP-SILL)^uwd^*^12^ larvae were collected in 1.5 ml microcentrifuge tubes. Embryo media was removed and larvae were incubated in 100 μL sterile, calcium-free ringer’s solution (116 mM NaCl, 2.6 mM KCl, 5 mM HEPES pH 7.0) at RT for 5 min. 600 μL of sterile protease solution (0.25% trypsin, 1 mM EDTA, pH 8.0, PBS) was warmed to 28°C and 27 μl of Collagenase P (100 mg/mL in HBSS) was added to the tube and mixed. Larvae were incubated at 28°C. Every 30s, larvae were homogenized for 30s by pipetting first with a P1000 pipette until partially dissociated, then with a P200 pipette until fully dissociated. Once dissociated, 100 μl of sterile stop solution (30% calf serum, 6 mM CaCl2, PBS) was added and mixed. The cells were spun down at 350xg at 4°C for 5 min. The supernatant was removed and replaced with 500 μl sterile suspension solution (1% FBS, 0.8 mM CaCl2, 50 U/mL penicillin, 0.05 mg/mL streptomycin, DMEM) chilled to 4°C. Cells were spun at 350xg at 4°C for 5 min. The supernatant was removed and 500 μl of chilled sterile suspension solution was used to resuspend cells. Resuspended cells were passed through a 40 μm cell strainer into FACS tube and stored on ice until sorting. Immediately prior to sorting, DAPI (1:1000) was added to sort out dead cells. GFP+ DAPI-cells were sorted on a BD FACSAria™ III Cell Sorter with a 100 μm nozzle into 500 μL TRIzol LS in a 1.5 mL microcentrifuge tube.

### Bulk RNA sequencing and analysis

Sorted pLL neurons were pooled into 3 replicates each for wild type and *actr10* samples. RNA was isolated using TRIzol. The SMART-Seq® v4 Ultra® Low Input RNA Kit was used for poly(A)+ selection, cDNA synthesis and amplification, followed by Illumina 150-bp paired-end sequencing performed by Azenta Life Sciences (South Plainfield, NJ). Sequence reads were trimmed to remove possible adapter sequences and nucleotides with poor quality using Trimmomatic v.0.36 ^105^. The trimmed reads were mapped to the Danio rerio GRCz10.89 reference genome available on ENSEMBL using the STAR aligner v.2.5.2b ^106^. Unique gene hit counts were calculated by using FeatureCounts from the Subread package v.1.5.2 ^107^. Only unique reads that fell within exon regions were counted. Using DESeq2 ^108^, a comparison of gene expression between WT and *actr10* samples was performed. The Wald test was used to generate p-values and log2 fold changes. Genes with an adjusted p-value < 0.05 and absolute log2 fold change > 1 were called as differentially expressed genes for each comparison. Alignment and differential expression analysis was performed by Azenta Life Sciences (South Plainfield, NJ).

KEGG pathway enrichment was performed for genes that were called as significantly upregulated or significantly downregulated using the ShinyGO (v0.82) webserver (*84-86*). *Danio rerio* gene IDs were used as input and background genes were defined as all genes that were detectable in the RNA-sequencing dataset. Default parameters were maintained. KEGG pathway plots were downloaded directly from the ShinyGO webserver. Gene Set Enrichment Analysis (GSEA) for differentially expressed genes was performed using the clusterProfiler R package with all gene ontology terms included, using the genome-wide annotation for Zebrafish (org.Dr.eg.db v3.19.1; ^109^). Zebrafish gene IDs were converted to their closest human gene orthologues using the g:Profiler g:Orth webserver ^110^. Fisher’s exact test enrichment analysis was performed using the fisher.test function in R to test overlap between significantly upregulated or significantly downregulated genes and either the MitoCarta database or the ESRRG ChipSeq targets ^49,63^. Transcription factor enrichment analysis was performed using the TFEA.ChIP webserver which uses information derived from the ChIP-seq datasets available from the ENCODE project, GEO Datasets, GeneHancer, ReMap 2018, and ReMap 2022 databases ^111^. The significantly downregulated genes were used with background genes defined as all genes that were detectable in the RNA-sequencing dataset. Wilcoxon rank-sum was used for transcription factor ranking. Over-representation analysis of transcription factor target motifs was performed using the Webgestalt webserver with the functional database identified as ‘‘network: transcription factor target” ^112^. The significantly downregulated genes were used with background genes defined as all genes that were detectable in the RNA-sequencing dataset. Default parameters were maintained. Volcano plots of differentially expressed genes and transcription factor enrichment were created using the ggplot2 R package ^113^.

### Experimental design, statistical analysis, and figure assembly

All image analysis was performed using ImageJ ^102^. Statistical analysis was performed using JMP18 and data plots made using GraphPad Prism 10, unless indicated otherwise in the method section. Sample size estimated based on previous work and determined by power analyses. Imaging and analysis done blind to genotype, allowing inherent randomization and blinding to experimental condition. All experiments were replicated no less than two times (experimental replicates) with sample size for each experiment representing biological replicates. Statistical tests/post hoc analyses and other statistical details for each data set are indicated in the text or figure legends. For parametric data, ANOVAs were used with Tukey HSD post-hoc contrasts for multiple pairwise comparisons within a dataset. For nonparametric data, single comparisons were done using Wilcoxon/Kruskal-Wallis analysis, multiple comparisons using the Steel-Dwaas test. Images were created using ImageJ and linear adjustments to brightness/contrast were made equally for images being compared. Figures were compiled in Adobe Illustrator and supplemental movies were handled in Adobe After Effects.

