## Supplemental information for "Retrograde mitochondrial transport is required for mitochondrial biogenesis in zebrafish neurons"

**Table S1.**

| Reagent or Resource | Source | Identifier | RRID |
|---|---|---|---|
| Antibodies | | | |
| Chicken anti-GFP | Aves | GFP-1020 | AB_10000240 |
| Rabbit anti-SSBP1 | ProteinTech | 12212-1-AP | AB_2195320 |
| Rabbit anti-TFAM | ProteinTech | 22586-1-AP | AB_11182588 |
| Sheep anti-DIG-AP | Roche | 11093274910 | AB_514497 |
| Drugs | | | |
| Resveratrol | MilliporeSigma | 554325 | |
| AICAR | Sigma-Aldrich | A9978 | |
| Experimental models: organisms/strains | | | |
| Zebrafish: AB | ZIRC | AB | ZDB-GENO-960809-7 |
| Zebrafish: *mitfa* (*nacre*) | [114] | w2 | ZDB-ALT-990423-22 |
| Zebrafish: *actr10* | [35] | nl15 | ZDB-FISH-170719-8 |
| Zebrafish: *p150a* | [37] | y625 | ZDB-ALT-200527-5 |
| Zebrafish: *p150b* | [35] | nl16 | ZDB-ALT-170628-4 |
| Zebrafish: *nudc* | [42] | nl21 | ZDB-ALT-190722-1 |
| Zebrafish: *TgBAC(neurod:eGFP)* | [115] | nl1 | ZDB-ALT-080701-1 |
| Zebrafish: *Tg(5kbneurod1:pgc1α-p2a-mRFP)* | [51] | uwd9 | ZDB-ALT-231026-11 |
| Zebrafish: *Tg(5kbneurod:mRFP-actr10)* | [37] | nl22 | ZDB-ALT-201006-6 |
| Zebrafish: *Tg(5kbneurod:mito-mEos)* | [37] | y586 | ZDB-ALT-180628-3 |
| Zebrafish*: Tg(hsp70l:EGFP-en.sill)* | This paper | uwd12 | ZDB-ALT-250520-12 |
| Recombinant DNA | | | |
| *5kbneurod1:mito-eGFP* | [51] | | |

| | |
|---|---|
| *5kbneurod1:mRFP* | [35] |
| *5kbneurod1:mitotagRFP* | [35] |
| *5kbneurod1:eGFP* | [51] |
| *Hsp701:Drp1K38A-mRFP* | [35] |
| *5kbneurod1:cox8a-cox8a-halotag* | This paper |
| *5kbneurod1:kif1a-egfp-omp25 ["MitoTruck"]* | This paper |
| *5kbneurod1:mRFP-esrra* | This paper |
| *5kbneurod1:mRFP-esrra*$^{4KR}$ | This paper |
| *5kbneurod1:cox8a-cox8a-paGFP-p2a-mitoTagRFP* | This paper |
| *5kbneurod:POLG2-GFP-p2a-mitoTagRFP* | This paper |
| *Hsp701:POLG2-GFP-p2a-mitoTagRFP* | This paper |
| *5kbneurod1:cox8a-cox8a-NAD-p2a-RFP* | This paper |
| *5kbneurod1:H2B-NAD-p2a-RFP* | This paper |
| *5kbneurod1:cox8a-cox8a-NAD-p2a-mitoTagRFP* | This paper |

Table S1: List of reagents and experimental models

**Table S2.**

| Primer | Sequence |
|---|---|
| Genotyping | |
| *actr10* FP | 5’- CTGTTTTCGGATGAACTGCCTG |
| *actr10* RP | 5’- ACTTACTTCGTGTAGGCCGC |
| *nudc* FP | 5’- GACCAGAGGCAGAAGTCCAT |
| *nudc* RP | 5’- TTTTTGGCAGCTTGTTTTGAAAG |
| *p150a* FP | 5’- TAGTGTGCAGATCCAATATGGC |
| *p150a* RP | 5’- ATTTCCCAGAGGCGAAGAGT |
| *p150b* FP | 5’- TCGGCTTCTGCAGGAGAGAT |
| *p150b* RP | 5’- CTGGGTCCCGGGATTGGAG |
| *in situ* probes | |
| *atp5md* FP | 5’- AATCATGGGTGGACACGACG |
| *atp5md* RP | 5’- AGCATGAAAATTGGTGCGGG |
| *cox5ab* FP | 5’- GATTTCAGCGGCGGTTCTTC |
| *cox5ab* RP | 5’- GAAGTCAAACTCGGTCCCGT |
| *nrf1* FP | 5’- ACACAAGTTGCCACAACAGC |
| *nrf1* RP | 5’- ACAGGTACTCCTTGGAGGGA |
| *pgc1a* FP | 5’- CTCGCAACATGGACGAAAGC |
| *pgc1a* RP | 5’- AATGGACACACTTTGCGCAG |
| *pgc1b* FP | 5’- AAAAGCGGATCTCAGCACCA |
| *pgc1b* RP | 5’- TTCGGCCATCCAGAGGATTG |
| *polb* FP | 5’- AACTGCGCAAGCTGGAAAAG |
| *polb* RP | 5’- GCCGTTGTGGATCATCTGGA |
| *tfam* FP | 5’- GGGGCTAATCTGCTGGTCAA |
| *tfam* RP | 5’- CGGCTTTTGCGGAGGAAAAA |
| *tomm40* FP | 5’- GGACATGGACCAACACCGAA |
| *tomm40* RP | 5’- GGGCCTAACCTATCAGGCTG |
| DNA plasmid generation | |
| *5kbneurod1:kif1a-egfp-omp25* ["MitoTruck"] | |
| *kif1a* FP | 5’- CTATAGGGCGAATTGGGTACATGGCCGGGGCCTCTGTG |
| *kif1a* RP | 5’- TGCTCACCATGGTAGGCCTAGGCGCGCC |
| *kif1a* GFP FP | 5’- TAGGCCTACCATGGTGAGCAAGGGCGGG |
| *kif1a* GFP RP | 5’- GGCCGTTCTGTCTAGAGACGGTCCGCTTGTAC |
| *omp25* MLS FP | 5’- CGTCTCTAGACAGAACGGCCCCACCAGC |
| *omp25* MLS RP | 5’- TATCAAGCTTATCGATACCGTTAAAATGGGCCCCGGGG |

*5kbneurod1:cox8a-cox8a-halotag*

| | |
|---|---|
| *cox8a_1* FP | 5’- CTATAGGGCGAATTGGGTACATGTCCGTCCTGACGCCG |
| *cox8a_1* RP | 5’- GGACGGACATTGGGTCCAACGAATGGATCTTG |
| *cox8a_2* FP | 5’- GTTGGACCCAATGTCCGTCCTGACGCCG |
| *cox8a_2* RP | 5’- TTTCGGATCCTGGGTCCAACGAATGGATCTTG |
| HaloTag FP | 5’- GTTGGACCCAGGATCCGAAATCGGTACTG |
| HaloTag RP | 5’- TATCAAGCTTATCGATACCGTTAACCGGAAATCTCCAG |

*hsp70l:EGFP-en.sill*

| | |
|---|---|
| Hsp70 GFP FP | 5’- TCCGCAGCCCCCAAGCTTGGATGGTGAGCAAGGGCGAG |
| Hsp70 GFP RP | 5’- GTTGGGATGGCTATAGGGCTGCAGAATCTAGAG |
| SILL FP | 5’- AGCCCTATAGCCATCCCAACTCACTCAC |
| SILL RP | 5’- TGGATCATCATCGATGGTACCTGACATTTTCCGGAACAG |

*5kbneurod1:esrra-p2a-mRFP*

| | |
|---|---|
| *esrra* FP | 5’- CTATAGGGCGAATTGGGTACATGTCTTCCAGAGAACGAC |
| *esrra* RP | 5’- CTTTGTACAAGGGTGAGTCCATCATGGC |
| p2a mRFP FP | 5’- GGACTCACCCTTGTACAAAGTGGGGGGATC |
| p2a mRFP RP | 5’- TATCAAGCTTATCGATACCGTTACTTGTACAAGGCGCC |

*5kbneurod1:cox8a-cox8a-NAD-p2a-RFP*

| | |
|---|---|
| mitoNAD FP | 5’- CTATGGGCGAATTGGGTACATGCTGGCCACCCGCGTG |
| mitoNAD RP | 5’- CTCCGGATCCACCTACACGTTGTGTCGGCG |
| RFP FP | 5’- ACGTGTAGGTGGATCCGGAGCCACGAAC |
| RFP RP | 5’- TATCAAGCTTATCGATACCGTTACTTGTACAAGGCGCCG |

*5kbneurod1:cox8a-cox8a-NAD-p2a-mitoTagRFP*

| | |
|---|---|
| mitomitoNAD FP | 5’-CTATAGGGCGAATTGGGTACGAATTGGGTACATGTCCG |
| mitomitoNAD RP | 5’- CTCCGGATCCACCTACACGTTGTGTCGG |
| p2a_mitoTag FP | 5’- ACGTGTAGGTGGATCCGGAGCCACGAAC |
| p2a_mitoTag RP | 5’- TATCAAGCTTATCGATACCGTTATAGTTTGTGCCCCAGTTTGC |

*5kbneurod1:H2B-NAD-p2a-RFP*

| | |
|---|---|
| H2B FP | 5’-CTATAGGGCGAATTGGGTACCGGGCCCCCCCTCGAGAC |
| H2B RP | 5’-CTTTTTCCATTCCCATGGTGGCGACCGG |
| NAD_p2a_RFP FP | 5’-CACCATGGGAATGGAAAAAGCTCCGCATG |
| NAD_p2a_RFP RP | 5’-TATCAAGCTTATCGATACCGTTACTTGTACAAGGCGCC |

*5kbneurod1:cox8a-cox8a-paGFP-p2a-mitoTagRFP*

| | |
|---|---|
| Cox8a1 FP | 5’-CTATAGGGCGAATTGGGTACATGTCCGTCCTGACGCCA |

| | |
|---|---|
| Cox8a1 RP | 5’-GGACGGACATCAACGAATGGATCTTGGCAC |
| Cox8aPAGFP FP | 5’-CCATTCGTTGATGTCCGTCCTGACGCCG |
| Cox8aPAGFP RP | 5’-CTCCGGATCCCTTGTACAGCTCGTCCATG |
| 2amitoTagRFP FP | 5’-GCTGTACAAGGGATCCGGAGCCACGAAC |
| 2amitoTagRFP RP | 5’- TATCAAGCTTATCGATACCGTTATAGTTTGTGCCCCAGTTTGC |
| *Hsp701:POLG2-GFP-p2a-mitoTagRFP* | |
| Polg2 FP | 5’-TCCGCAGCCCCCAAGCTTGGATGCGCTCTCGTGTAGCC |
| Polg2 RP | 5’- TGGATCATCATCGATGGTACTTATAGTTTGTGCCCCAGTTTGC |
| *5kbneurod1:mRFP-esrra* | |
| mRFP FP | 5’- CTATAGGGCGAATTGGGTACATGTCTTCCAGAGAACGAC |
| mRFP RP | 5’- TGGAAGACATGTACAAGGCGCCGGTGGA |
| *esrra* FP | 5’- CGCCTTGTACATGTCTTCCAGAGAACGAC |
| *esrra* RP | 5’ - TATCAAGCTTATCGATACCGCTAGGGTGAGTCCATCATG |
| Mutagenesis | |
| *5kbneurod1:mRFP-esrra*$^{4KR}$ | |
| aa129 FP | 5’- CCAAGCGCCGCAGAAGGGCTTGTCAGG |
| aa129 RP | 5’- CCTGACAAGCCCTTCTGCGGCGCTTGG |
| aa138 FP | 5’-CGTGTCGCTTCACCAGGTGCCTCAAAGTAGG |
| aa138 RP | 5’- CCTACTTTGAGGCACCTGGTGAAGCGACACG |
| aa160_162 FP | 5’- GGTGGACGACAGAGGTACAGGAGACGCCCTGAG |
| aa160_162 RP | 5’- CTCAGGGCGTCTCCTGTACCTCTGTCGTCCACC |

Table S2: List of oligonucleotide primers

**Table S3.**

| Gene | ENSEMBL ID | Probe set size, amplifier |
|---|---|---|
| *atf3* | ENSDART00000022060.8 | 16, B2 |
| *atp5md* | ENSDART00000150789.3 | 5, B1 |
| *cox5ab* | ENSDART00000172518.2 | 10, B1 |
| *cox6c* | ENSDART00000056319.4 | 6, B1 |
| *cox7c* | ENSDART00000165249.2 | 5, B2 |
| *cox8a* | ENSDART00000138236.2 | 8, B1 |
| *efhd1* | ENSDART00000063779.4 | 8, B1 |
| *ndufa5* | ENSDART00000103293.4 | 7, B2 |
| *ndufb3* | ENSDART00000144625.3 | 7, B2 |
| *pgc1a* | ENSDART00000097710.6 | 20, B1 |
| *polb* | ENSDART00000002764.9 | 20, B1 |
| *spry4* | ENSDART00000099528.4 | 18, B2 |
| *tfam* | ENSDART00000092009.6 | 20, B2 |
| *tomm40* | ENSDART00000015302.7 | 18, B2 |
| *uqcrq* | ENSDART00000180722.1 | 5, B2 |

Table S3: List of probe sets used for HCR RNA FISH

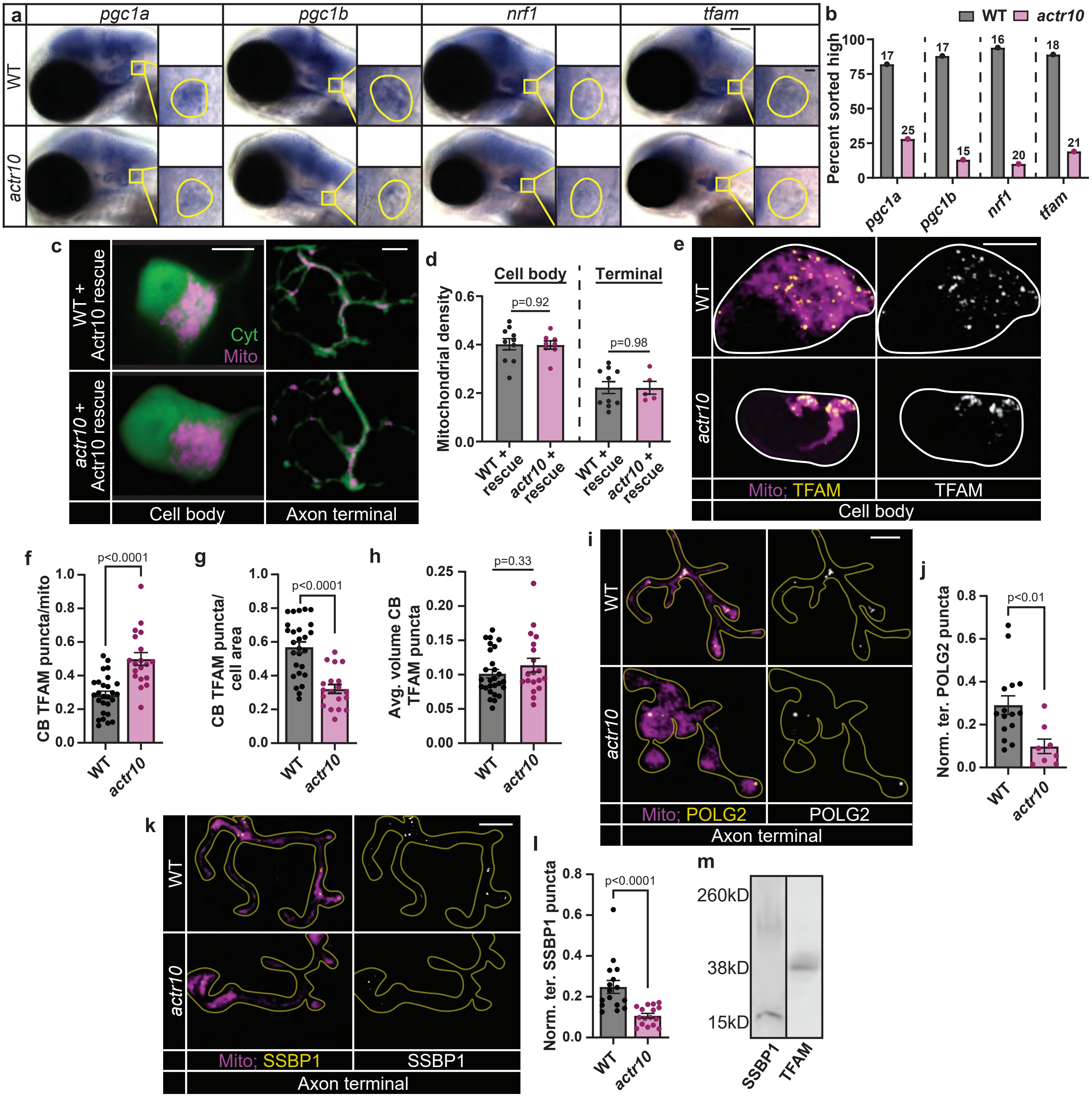

**Supplementary Figure 1: Mitochondrial biogenesis markers are reduced in *actr10* mutant neurons.** (a) Digoxigenin-labeled in situ hybridization for *pgc1a, pgc1b, nrf1*, and *tfam* mRNA in 4 dpf zebrafish. Inset shows staining in the pLLg (outline). (b) Larvae were sorted into high or low mRNA expression in the pLLg or brain and genotyped post-sorting. Percentage of total wild type or *actr10* larvae sorted "high expression" is shown. Number of larvae per group indicated on graph. (c) 4 dpf pLL single cell body and axon terminal for wild type and *actr10* mutant larvae expressing Actr10 in neurons (+ Actr10 rescue; *Tg(neurod:mRFP-Actr10)*$^{nl15}$). Cytosolic area visualized with GFP (Cyt); mitochondria visualized with matrix-localized HaloTag (Mito, magenta). (d) Quantification of mitochondrial density (mitochondrial area/cytosolic area; ANOVAs). (e) TFAM immunostaining in pLL cell body mitochondria, visualized with mitochondria-localized GFP (Mito, magenta; cell outlined). (f) Number of TFAM puncta normalized to mitochondrial volume (Wilcoxon). (g) Number of TFAM puncta normalized to cell body area (ANOVA). (h) Average volume of TFAM puncta (Wilcoxon). (i) pLL axon terminals expressing POLG2-GFP and mitoTagRFP (Mito, magenta; terminal outlined). (k) SSBP1 immunostaining in pLL axon terminal mitochondria, visualized with mitochondria-localized GFP (Mito, magenta; terminal outlined). (j, l) Number of POLG2 or SSBP1 puncta normalized to mitochondrial volume (Wilcoxon). (m) Anti-SSBP1 and Anti-TFAM western blots of wild type zebrafish larval protein extracts. Scale bars: (a) Head = 100 µm, inset = 10 µm; (c, e, i, k) = 5 µm. All data are mean ± SEM. Data points represent individual larvae.

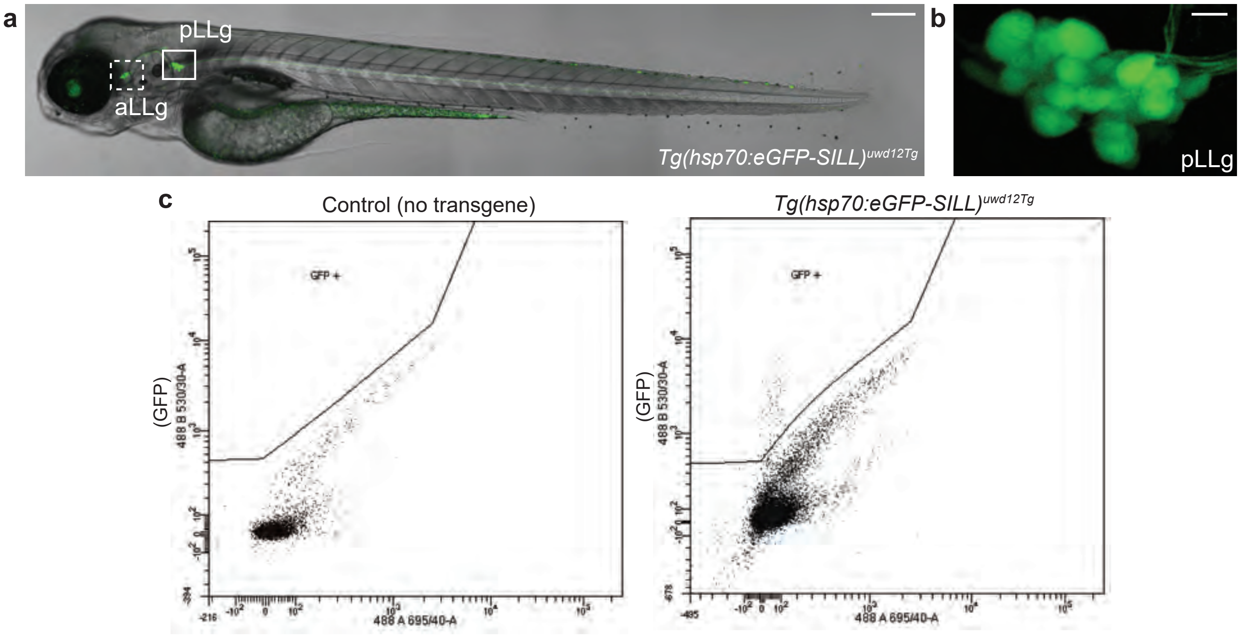

**d** Gene Set Enrichment Analysis of Gene Ontology

| Description | Set Size | Enrichment Score (ES) | Normalized ES | p.adjust |
|---|---|---|---|---|
| mitochondrial protein-containing complex | 210 | -0.5326 | -2.28005 | 4.54E-08 |
| inner mitochondrial membrane protein complex | 131 | -0.56575 | -2.29566 | 1.79E-06 |
| mitochondrial inner membrane | 275 | -0.4498 | -1.98734 | 8.85E-06 |
| ribosome | 178 | -0.49632 | -2.07555 | 1.30E-05 |
| structural constituent of ribosome | 150 | -0.51605 | -2.12838 | 3.11E-05 |
| ribosomal subunit | 144 | -0.51706 | -2.11537 | 4.92E-05 |
| organelle inner membrane | 292 | -0.42636 | -1.89981 | 4.92E-05 |
| mitochondrial respiratory chain complex I | 39 | -0.73818 | -2.40085 | 5.10E-05 |
| NADH dehydrogenase complex | 39 | -0.73818 | -2.40085 | 5.10E-05 |
| respiratory chain complex I | 39 | -0.73818 | -2.40085 | 5.10E-05 |

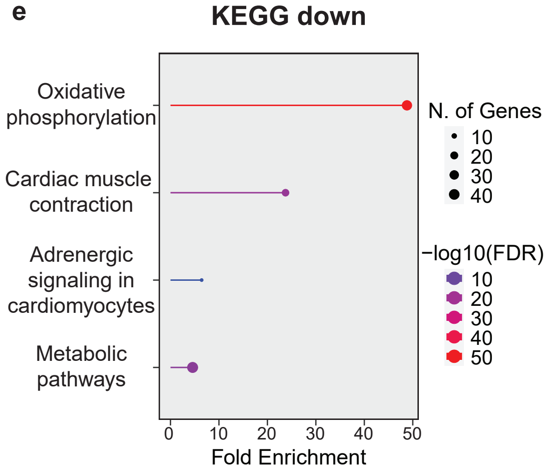

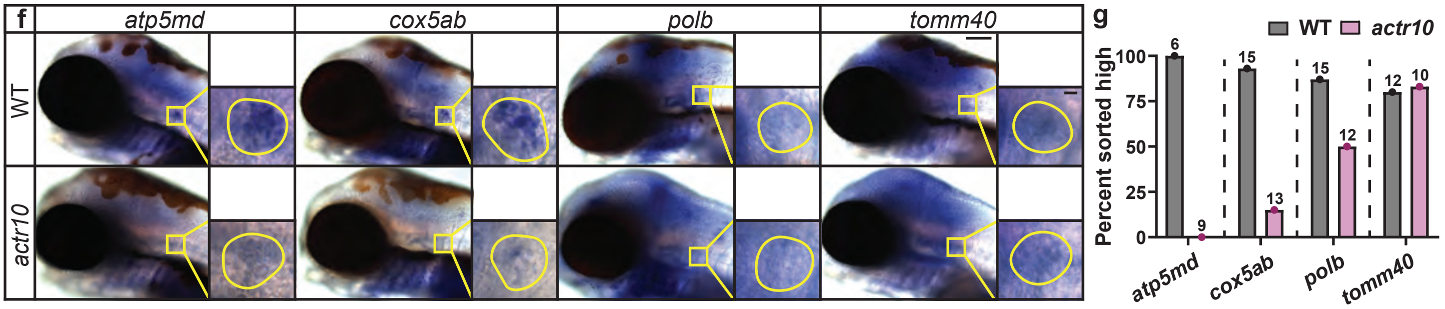

**Supplementary Figure 2: Mitochondrial gene transcription is impaired in *actr10* mutants.** (a) 4 dpf zebrafish transgenic larva carrying the *Tg(hsp70:eGFP-SILL)*$^{uwd12Tg}$ transgene used for FACS. GFP expression in the anterior (aLLg) and posterior (pLLg) lateral line ganglions is indicated. Weak green in eye, yolk, and pigment cells is due to autofluorescence and was excluded using optimized FACS gating parameters. Scale bar = 200 µm. (b) Magnified pLLg from *Tg(hsp70:eGFP-SILL)*$^{uwd12Tg}$ transgenic zebrafish in (a). Scale bar = 10 µm. (c) FACS gating parameters used to isolate true GFP+ cells from autofluorescent cells. (d) Gene Set Enrichment Analysis results for the top 10 de-enriched gene ontology terms in *actr10* mutants relative to wild type. (e) Enrichment analysis of KEGG pathways for significantly downregulated genes in *actr10* mutants. (f) Digoxigenin-labeled in situ hybridization for *atp5md, cox5ab, polb,* and *tomm40* mRNA in 4 dpf larvae. Inset shows staining in the pLLg (outlined). Scale bars: Head = 100 µm, inset = 10 µm. (g) Larvae were sorted into high or low mRNA expression in the pLLg or brain and genotyped post-sorting. Percentage of total wild type or *actr10* larvae sorted “high expression” is shown. Number of larvae per group indicated on graph.

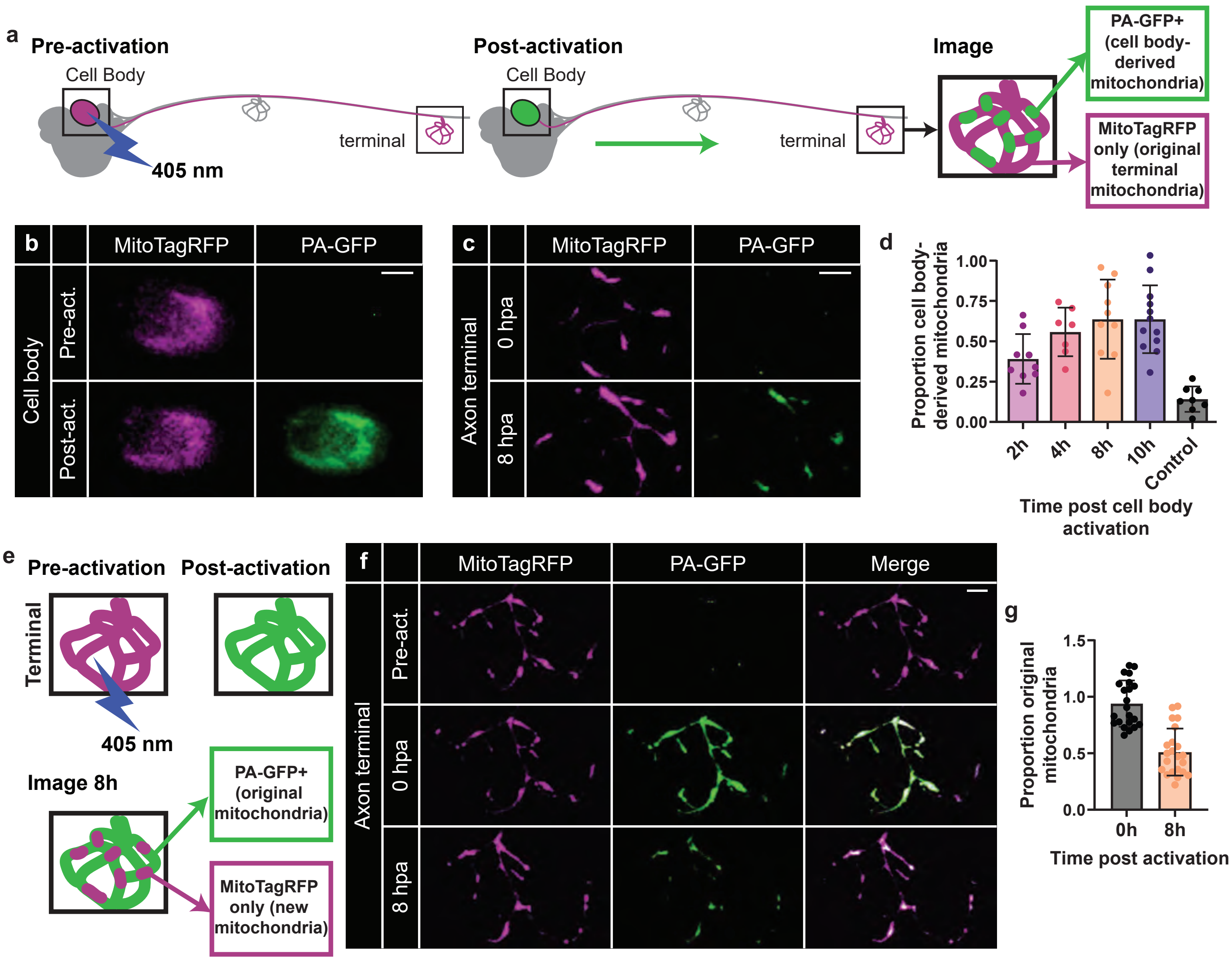

**Supplementary Figure 3: Cell body-derived mitochondria contribute significantly to the axon terminal.** (a) pLL cell body activation strategy to quantify cell body-derived mitochondria that contribute to the axon terminal. A pLL cell body co-expressing MitoTagRFP and mitochondrial matrix-localized photoactivateable (PA) GFP was locally activated using 405 nm laser and the corresponding axon terminal was imaged at set time points post cell body activation (hpa). (b) A pLL cell body pre- and post-activation (act.). (c) A pLL axon terminal 0 hpa and 8 hpa. (d) Proportion of cell body derived (PA-GFP+) mitochondria in the axon terminal relative to all axon terminal mitochondria (MitoTagRFP+) at multiple time points post cell body activation. Control was an axon terminal imaged at 0h and 8h with no cell body activation. (e) Terminal photoactivation strategy to assess mitochondrial turnover. A pLL axon terminal co-expressing MitoTagRFP and mitochondrial matrix-localized PA-GFP was locally activated using 405 nm laser and imaged 8 hpa. (f) A pLL axon terminal pre-activation, immediately post-activation (0 hpa) and 8 hpa. (g) Proportion of original (PA-GFP+) mitochondria in the axon terminal relative to all mitochondria (MitoTagRFP+). Scale bars = 5 µm. All data are mean ± SEM; data points represent individual larvae.

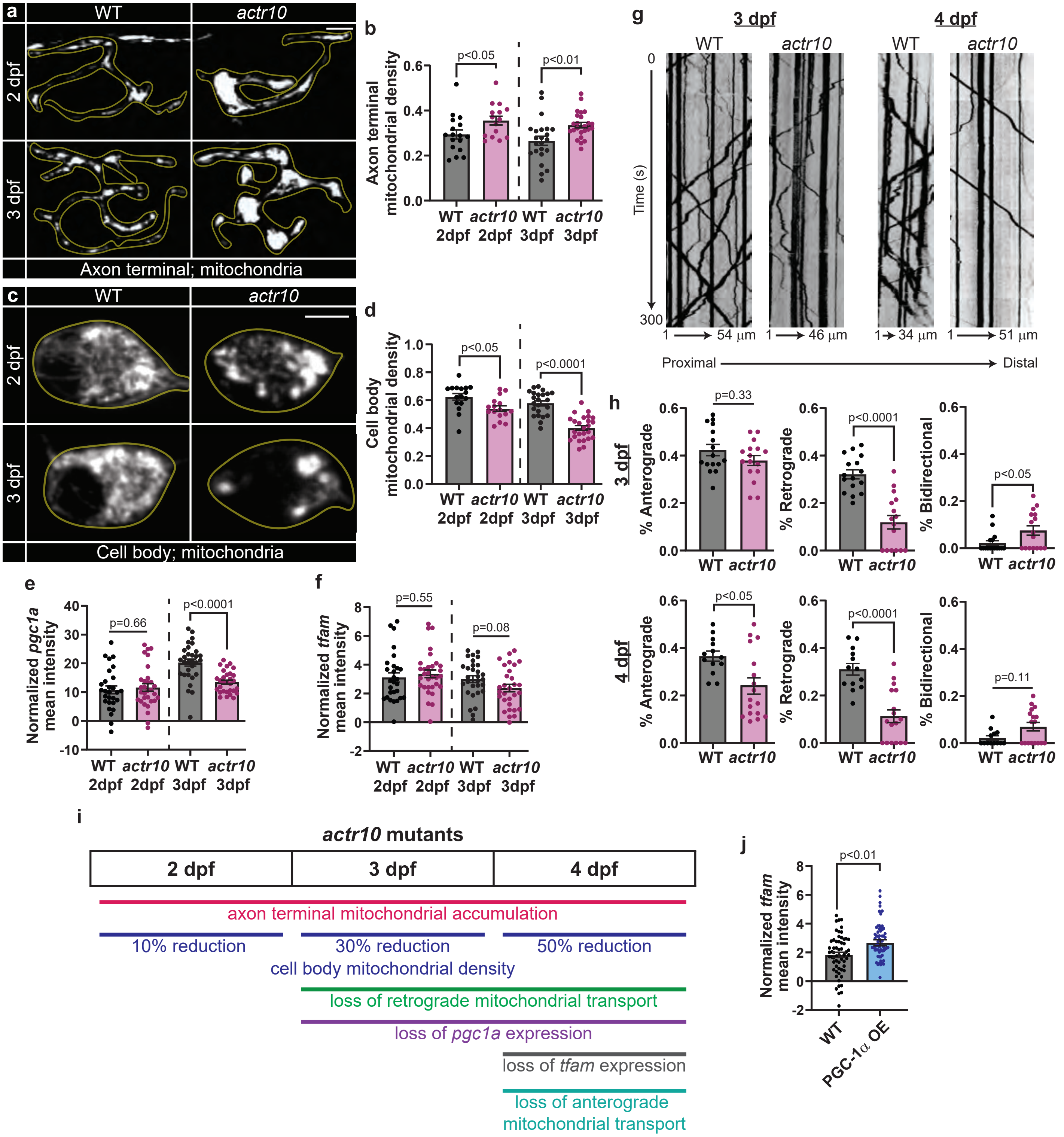

**Supplementary Figure 4: *actr10* mutants display progressive loss of mitochondrial biogenesis.** (a, c) pLL axon terminals or cell bodies (outlined) expressing mitoTagRFP in 2 and 3 dpf wild type or *actr10* mutant larvae. (b, d) Quantification of mitochondrial density (mitochondrial area/cytosolic area; ANOVAs). (e, f) Mean fluorescence intensity of *pgc1a* (e) and *tfam* (f) HCR RNA FISH normalized to background in the pLLg of 2 and 3 dpf wild type and *actr10* mutant larvae (ANOVAs). (g) Representative kymographs of mitochondrial transport in wild type and *actr10* pLL axons at 3 and 4 dpf. (h) Mitochondrial transport frequencies (Wilcoxons). Frequencies represent the percentage of mitochondria moving anterogradely, retrogradely, or bidirectionally out of total mitochondria in the region imaged. (i) Summary of changes to mitochondrial localization and transport relative to mitochondrial biogenesis measures over time in *actr10* mutants compared to wild type controls. (j) pLLg mean fluorescence intensity of *tfam* HCR RNA FISH normalized to background for wild type vs. PGC-1α overexpression (OE; ANOVA). Scale bars = 5 µm. All data are mean ± SEM and data points represent individual larvae.

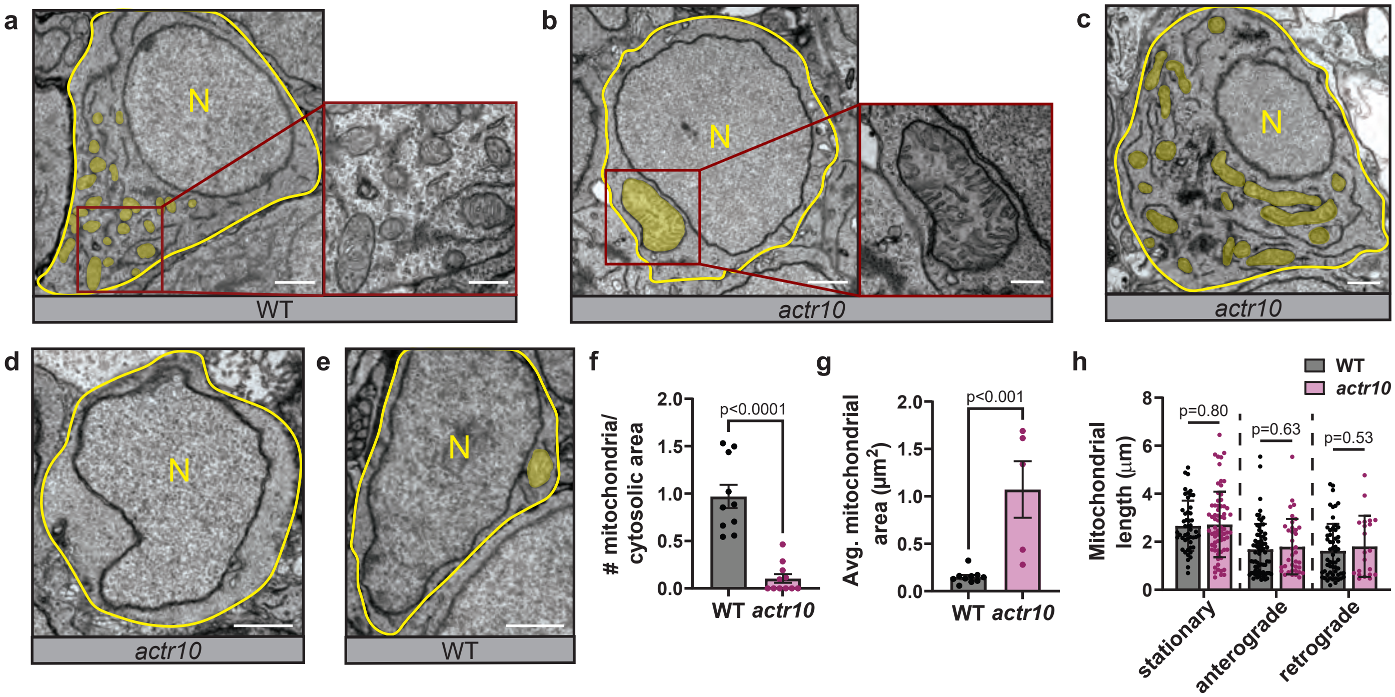

**Supplementary Figure 5: Mitochondrial structure in *actr10* mutants.** (a-e) TEM images of pLL cell bodies. Cell area outlined in yellow, nucleus marked with "N," and mitochondria shaded in yellow. Scale bars: Full cell = 1 µm, insets = 400 nm. (f, g) Quantification of number of mitochondria relative to cytosolic area (cell area - nuclear area) and average mitochondrial area from TEM images (Wilcoxons). TEM data points represent individual cell bodies from one ganglion per genotype. (h) Length of stationary and motile mitochondria in wild type vs. *actr10* mutant pLL axons at 4 dpf. Data points represent all mitochondria measured from 10 wild type and 15 *actr10* mutant axons (ANOVAs). All data are mean ± SEM.

**a** TFEA.ChIP Transcription factor enrichment of downregulated genes

| TF | Cell Type | Treatment | log2(OR) | log10(adj. p-value) | Euclidean Distance |
|---|---|---|---|---|---|
| ESR1 | MCF-7 | ethanol | 2.8140 | 1.6742 | 6.2604 |
| ZNF527 | HEK293T | N/A | 2.2369 | 1.2989 | 3.9344 |
| ESRRG | BT-474 | N/A | 1.4826 | 3.3718 | 3.8196 |
| MED12 | MCF-7 | CTL | 2.0359 | 1.7679 | 3.5692 |
| ESRRA | BT-474 | N/A | 1.3432 | 3.0862 | 3.4478 |
| SOX2 | NPC | R1159Q | 2.0844 | 0.9761 | 3.3848 |
| EZH2 | myotube | N/A | 1.2964 | 2.9988 | 3.3336 |
| TCF3 | Jurkat | N/A | 2.0275 | 0.7168 | 3.1594 |
| TP53 | MOLM-13 | M237I Daunorubicin | 1.9877 | 0.9321 | 3.1091 |
| TP53 | MOLM-13 | Y220C Daunorubicin | 1.9877 | 0.9321 | 3.1091 |
| EZH2 | A-673 | N/A | 1.2510 | 2.7131 | 3.0440 |
| EZH2 | HUVEC | VEGF 12h | 1.2808 | 2.6658 | 3.0250 |
| EP300 | HCT-116 | Nutlin3a | 1.8991 | 1.2757 | 3.0133 |
| REST | neuroblastoma | SKNMM | 1.7623 | 1.7851 | 2.9849 |
| YAP1 | HUCCT1 | N/A | 1.9569 | 0.6898 | 2.9636 |
| EZH2 | LNCaP | N/A | 1.1427 | 2.6407 | 2.9039 |
| SUZ12 | HEK293T | PCGF1352fl | 1.3262 | 2.4649 | 2.8892 |
| TP53 | MCF-7 | NUT | 1.8560 | 1.0619 | 2.8271 |
| ZNF774 | HEK293 | N/A | 1.8896 | 0.6657 | 2.7861 |
| REST | NCI-H295R | SF1 | 1.6389 | 1.7685 | 2.7564 |
| EZH2 | hepatocyte | N/A | 1.0598 | 2.3325 | 2.5724 |
| ZNF274 | GM08714 | N/A | 1.8178 | 0.3966 | 2.5564 |
| AR | DU145 | ARG56W | 1.7708 | 0.8293 | 2.5510 |
| EZH2 | fibroblast | dermal | 1.0443 | 2.1829 | 2.4277 |
| EZH2 | VCaP | DHAT 2H | 1.2190 | 2.0300 | 2.4257 |
| EZH2 | THP-1 | N/A | 0.9693 | 2.2256 | 2.4230 |
| ESRRA | BT-474 | AICAR | 0.9514 | 2.2304 | 2.4180 |
| JARID2 | UtE-iPS-7 | N/A | 1.0166 | 2.1829 | 2.4108 |
| EPAS1 | HUVEC | 16h 1%O2 | 1.7337 | 0.6140 | 2.4054 |
| NOTCH1_NICD | REC-1 | GSI-mock-washout | 1.7337 | 0.6140 | 2.4054 |

**b** Transcription factor target enrichment of downregulated genes

| Gene Set | Set Size | Enrichment Ratio | p-value | FDR |
|---|---|---|---|---|
| TGACCTY_V$ERR1_Q2 | 755 | 3.0122 | 1.99E-07 | 6.55E-05 |
| TGACCTTG_V$SF1_Q6 | 190 | 5.9847 | 2.14E-07 | 6.55E-06 |
| TGACATY_UNKNOWN | 475 | 2.578 | 0.00094 | 0.1656 |
| GCANCTGNY_V$MYOD_Q6 | 655 | 2.2702 | 0.00109 | 0.1656 |
| V$AMEF2_Q6 | 192 | 3.6446 | 0.00154 | 0.1656 |
| V$NKX61_01 | 156 | 3.9249 | 0.00200 | 0.1656 |
| V$SF1_Q6 | 202 | 3.4641 | 0.00212 | 0.1656 |
| YTATTTTNR_V$MEF2_02 | 518 | 2.364 | 0.00216 | 0.1656 |
| V$MMEF2_Q6 | 208 | 3.3642 | 0.00254 | 0.17355 |
| V$OCT1_05 | 176 | 3.4789 | 0.00393 | 0.22473 |

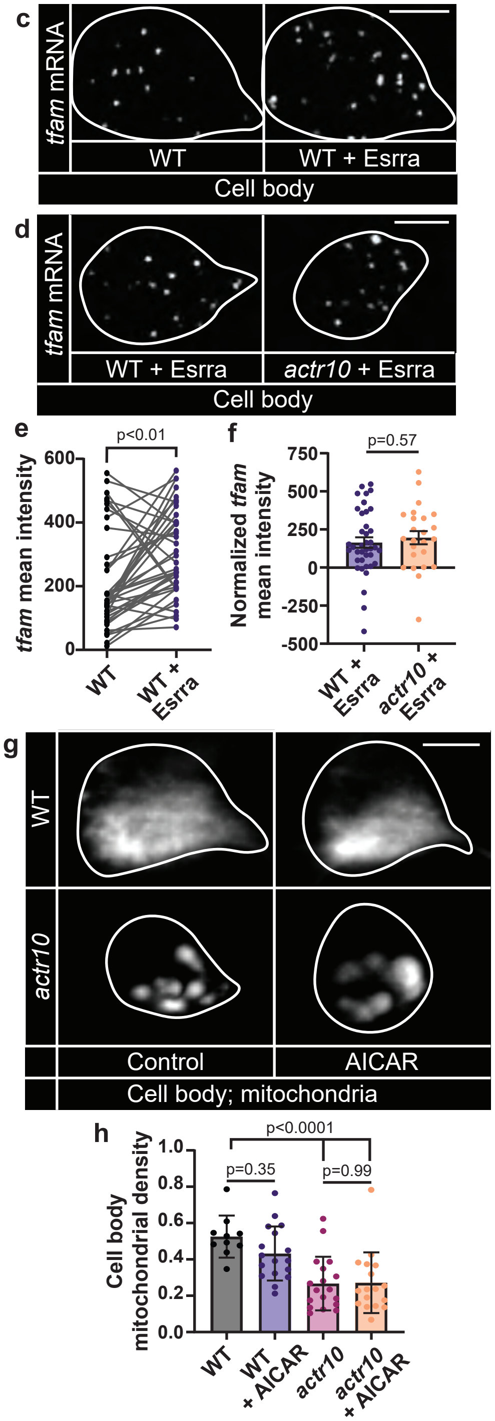

**Supplementary Figure 6: Estrogen related receptors mediate the transcriptional regulation of mitochondrial biogenesis.** (a) Transcription factor enrichment analysis for genes significantly downregulated in *actr10* mutants using TFEA.ChIP. Top 30 transcription factors (TF) based on euclidean distance from log2(Odds Ratio)/log10(adjusted p-value) to 0 are listed. (b) Transcription factor target enrichment analysis showing consensus motifs in the promoters of significantly downregulated genes. (c) HCR RNA FISH labeling of *tfam* mRNA in single pLL cell bodies (outlined) expressing Esrra or an adjacent control cell body in the same pLLg (WT). (d) HCR RNA FISH labeling of *tfam* in a single pLL cell body (outlined) of a wild type and *actr10* larva expressing Esrra. (e) Mean fluorescence intensity of *tfam* (paired t-test). (f) Mean fluorescence intensity of *tfam* normalized to background (ANOVA). (g) Mitochondria in wild type and *actr10* pLL cell bodies (outlined) with and without AICAR treatment. (h) Quantification of mitochondrial density (mitochondrial area/cytosolic area; ANOVAs with Tukey HSD). Data points represent individual larvae; data are mean ± SEM. Scale bars = 5 µm.
